# Adaptation of *Anaplasma phagocytophilum* to the tick vector is controlled by the transcriptional regulator Tr1

**DOI:** 10.1101/2025.11.12.688119

**Authors:** EricaRose Warwick, Rachel Burt, Jeffrey T. Badigian, Daniel Howell, Kyle T. Swallow, Chloe Leach, Azeza M. Falghoush, Dana K. Shaw, Ian T. Cadby, Jason M. Park

## Abstract

Rickettsial pathogens are strictly dependent on the cellular biology of their hosts for survival and replication. Predominantly transmitted by blood-feeding arthropods, these vector-borne pathogens are forced to adapt between the disparate environments of their mammalian host and arthropod vector. To achieve this, the Rickettsial bacteria *Anaplasma phagocytophilum* undergoes extensive transcriptional reprogramming with over 41% of its genes differentially transcribed between mammals and *Ixodes scapularis* ticks. How the bacterium orchestrates this dramatic transcriptional reprogramming is not understood. The gene *tr1* encodes a Helix-Turn-Helix DNA-binding protein that is exclusively expressed during tick infection. Herein, we show that *tr1* is essential for *A. phagocytophilum* survival in ticks and regulates the transcription of other genes necessary to adapt to the arthropod vector. We demonstrate that Tr1 is a DNA-binding protein and recognizes promotors of tick-specific genes in *A. phagocytophilum,* including secreted effector *ateA*, alternate components of type IV secretion system (T4SS), and membrane proteins. Our findings demonstrate that Tr1 is a master regulator of genes that are critical for *A. phagocytophilum* adaptation to the tick.

## INTRODUCTION

Rickettsial pathogens are strictly dependent on the cellular biology of their hosts for survival and replication, which is reflected by their small genomes and reduced metabolic capacities. Instead these bacteria have evolved factors to mediate host-pathogen interactions and manipulate the intracellular environment^1–4^. These include specialized surface proteins mediating contact and uptake into cells^2,3,5–9^ and effector molecules injected through secretion systems to redirect host cell pathways^10–23^. The large majority of rickettsial host-pathogen interactions are described in the context of mammalian infection^17,20–22,24,25^. However, since rickettsial bacteria are predominantly transmitted by blood-feeding arthropods, these findings only explore half of the pathogen lifecycle^26^. Mammals and arthropods are separated by over 680 million years of evolution and, as such, present distinct environments that Rickettsial pathogens must adapt to survive intracellularly ^27^.

The most common rickettsial pathogen in the United States, *Anaplasma phagocytophilum*, is transmitted to humans, domestic animals, and wildlife by the tick *Ixodes scapularis*^28^. While in the mammalian host, *A. phagocytophilum* infects and replicates within circulating neutrophils. In the tick, the bacteria initially infect the digestive tract and then migrate to the salivary glands, where they will survive through the molt^29,30^. In response to the disparate biology between the mammalian host and tick vector^31^, *A. phagocytophilum* undergoes extensive transcriptional reprogramming. Transcriptomic studies found that over 41% of *A. phagocytophilum* genes are differentially transcribed when comparing infected human monocytes and tick cells^32,33^. Indeed, Himar1 transposon disruption of host or vector specific *A. phagocytophilum* genes reduces bacterial survival in the respective human or tick cell cultures^10,34–37^. Together, this suggests transcriptional reprogramming of *A. phagocytophilum* is necessary for adaptation between the host and vector environments.

While both transcriptomics and proteomics have shown *A. phagocytophilum* and other rickettsial pathogens undergo extensive retooling during tick infections^5,32,33,38–42^, the regulatory mechanisms that control these dramatic shifts are not known. One gene, *tr1*, encodes a putative Helix-Turn-Helix DNA-binding protein (Tr1). *tr1* was first noticed due to its proximity to outer membrane protein (OMP) genes *omp_1X*, *omp_1N* and the *msp2* expression site^43,44^, but its impact on this neighboring operon is not defined. Strikingly, of all *A. phagocytophilum* genes *tr1* has the highest transcriptional specificity for tick cell infection^32,33^. Herein, we demonstrate that *tr1* is essential for *A. phagocytophilum* survival in tick cells, colonization of ticks *in vivo*, and expression of many genes necessary for adaptation to the tick. Further, we demonstrate that Tr1 is a DNA-binding protein and recognizes promotors of tick-specific genes in *A. phagocytophilum,* including secreted effector *ateA*, alternate components of type IV secretion system (T4SS), and membrane proteins. From this work we have identified Tr1 as a master regulator of genes that are critical for *A. phagocytophilum* adaptation to the tick.

## RESULTS

### Transcription of tr1 is specific to tick cell infection

*A. phagocytophilum* HGE1 *tr1* transcription was quantified and compared between bacteria cultured in the human monocyte-like HL60 cell line and embryonic tick ISE6 cells. Consistent with previous tiling array reports, *tr1* transcription was >26 fold higher during growth in tick ISE6 cells over the human HL60 cell line (**Fig 1a**), validating that *tr1* expression is highly specific to growth in tick cells^32,33^.

**Figure 1.**
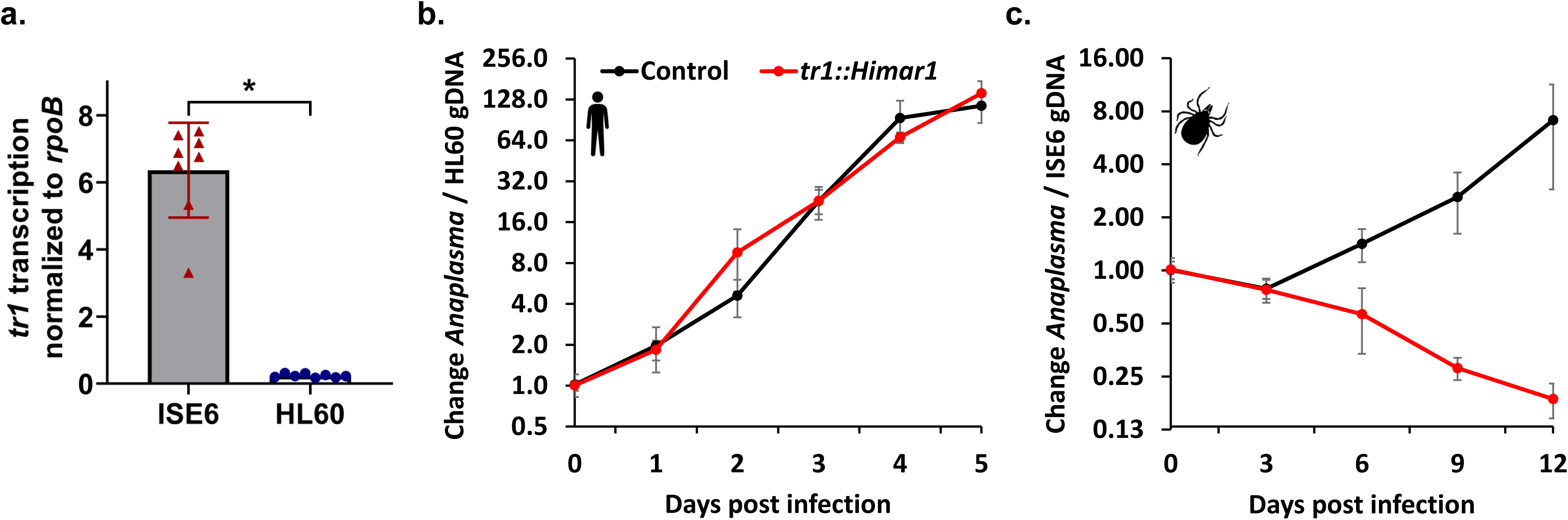
*tr1* transcription is specific to tick cell infection and necessary for *A. phagocytophilum* survival in tick cells. (**a**) *tr1* transcription normalized to housekeeping gene *rpoB* during *A. phagocytophilum* culture in ISE6 tick cells and human HL60 cells. Bar indicate mean of four replicate infections each measured in two technical replicates show as points. (**b and c**) Growth of *A. phagocytophilum tr1*::Himar1 or control strain in cell culture infections of (**b**) human HL60 cells and (**c**) tick ISE6 cells. *A. phagocytophilum* burden measured by bacterial gDNA relative to eukaryotic host gDNA via qPCR. Data displayed as mean with ±SD of three biological replicates with two technical replicates each. Data are representative of three experimental replicates. \**P* < 0.05 (Mann-Whitney *t*-test).

### tr1 is necessary for A. phagocytophilum survival in tick cells

The specificity of *tr1* transcription to ISE6 cell culture prompted us to test if *tr1* is necessary for *A. phagocytophilum* adaptation to ticks. To examine this possibility, we obtained a transposon mutant strain *tr1*::Himar1 from the previously published mutant collection^34^. An established control strain with the Himar1 transposon inserted in an intergenic non-coding region was used for comparison. This control strain is phenotypically comparable to wild-type in both mammalian and tick infection models^10,34,36,37^. *tr1::Himar1* and the control *A. phagocytophilum* strain were purified from HL60 cell culture and used to infect either ISE6s or HL60s. Bacterial burdens were compared over time. During HL60 infection, the control and *tr1*::Himar1 strains grew equivalently (**Fig 1b**). However, during tick cell infection, *tr1*::Himar1 steadily declined (**Fig 1c**), indicating that it is required for *A. phagocytophilum* adaptation to tick cells.

### A. phagocytophilum requires tr1 to colonize ticks in vivo

The *in vitro tr1*::Himar1 phenotype led us to ask if *tr1* is similarly required *in vivo* for *A. phagocytophilum* adaptation to the tick. Mice were infected with either the intergenic control or the *tr1*::Himar1 mutant *A. phagocytophilum*. Seven days post infection bacterial burden in the mice was quantified and found to be equivalent between the strains, indicating *tr1* is not required during murine infection (**Fig 2a**). Burden matched mice from these groups were used to feed *I. scapularis* larval ticks to repletion. We found ticks that fed on mice infected with *tr1*::Himar1 acquired significantly less *A. phagocytophilum* than those that fed on mice infected with the control strain (Fig 2b). These findings indicate that *tr1* is dispensable for murine infection but required for colonization of the arthropod vector.

**Figure 2.**
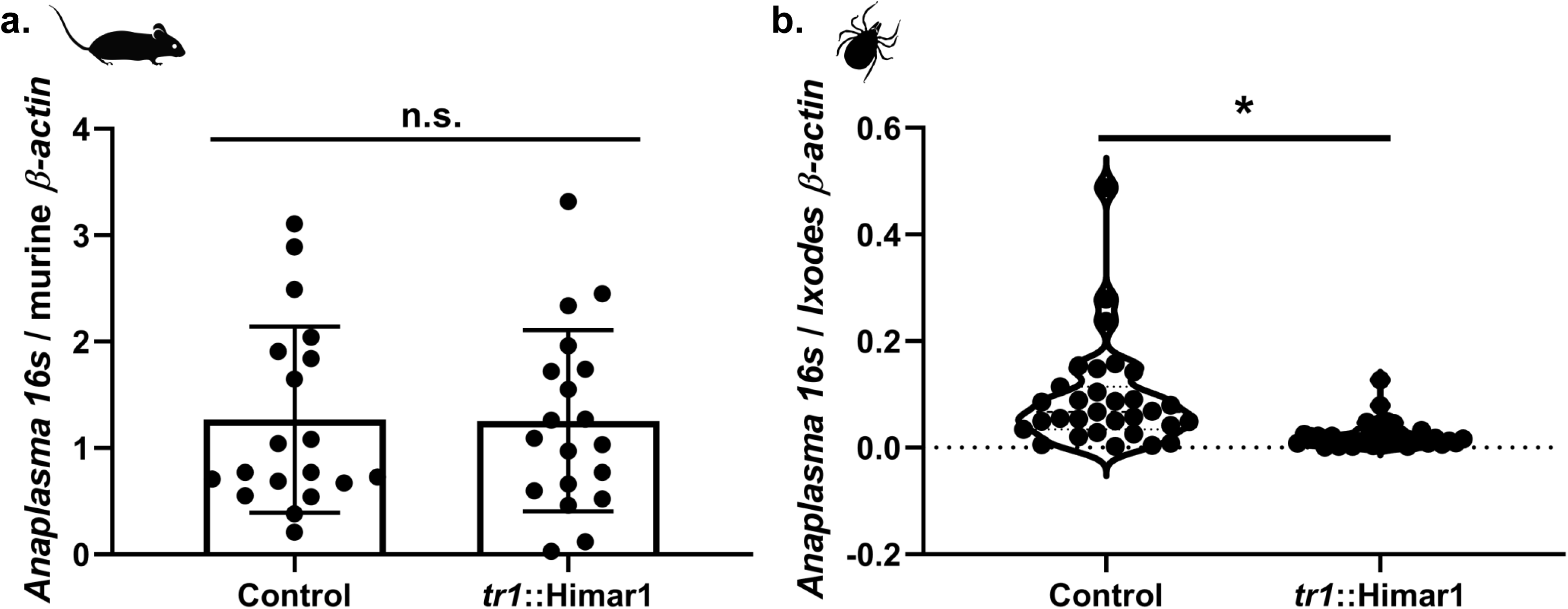
*tr1* is despicable during murine infection but essential for acquisition by ticks. (**a**) Mouse blood *Anaplasma* burden 7 days post intraperitoneal inoculation with 1 × 10^8^ *A. phagocytophilum tr1*::Himar1 or control::Himar1 strains. Blood processed for gDNA and bacterial burden was measured by qPCR of *A. phagocytophilum* 16S rDNA versus mouse actin by ΔΔCt. Each strain was tested in twenty mice (½ male, ½ female). Each sample was tested in technical duplicate reaction. (**b**) Two burden-matched infected mouse pairs were used for *Ixodes scapularis* larvae infestation. Ticks were allowed to feed to repletion and detach. Whole replete *I. scapularis* larvae were processed for RNA. *A. phagocytophilum* bacterial loads were measured via qRT-PCR of *A. phagocytophilum* 16S rRNA levels versus mouse actin transcripts. Data includes ticks from two burden matched mouse pairs. From each mouse, 10–20 individual ticks were collected as biological replicates, and each qRT-PCR was performed in duplicate. \**P* < 0.005 (Welch’s *t*-test).

### Tr1 is a multimeric Helix-Turn-Helix DNA-binding protein

Having established that Tr1 is important for adaptation to the tick host by *A. phagocytophilum*, we next used bioinformatics approaches to predict the functions of the Tr1 protein. First, a structural model of monomeric Tr1 protein (amino acids 1-183) was generated using ColabFold^45^. The resulting predicted Tr1 monomer model is comprised of two ordered and predominantly α-helical domains that are separated by a central disordered peptide linker and flanked by disordered peptide at both the N and C termini (**Fig 3a**). In total, the model contains approximately 58 residues of predicted disorder. Whilst the global confidence score for the predicted model was low (pTM=0.468), the local confidence scores for the two ordered domains were relatively high (pLDDT greater than 80 for residues ∼25-90 and ∼130-155, Supplementary figure 1).

**Figure 3.**
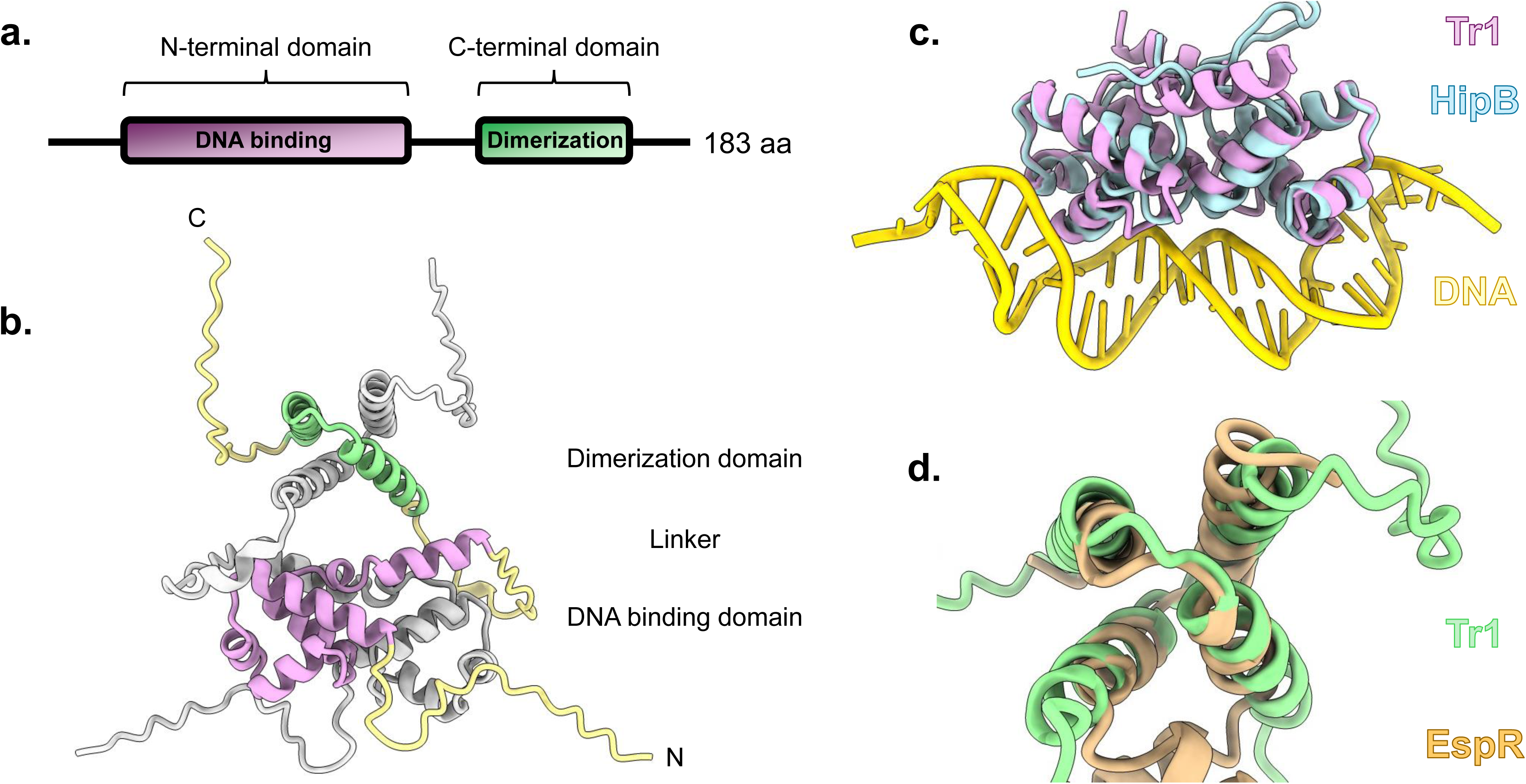
Structural modelling of the Tr1 protein. **(a)** Schematic of the predicted Tr1 protein domain architecture. Ordered domains are displayed as boxes and predicted disordered regions as solid black lines. **(b)** Cartoon representation of a predictive Tr1 dimer protein model. For one Tr1 chain the N-terminal DNA binding domain is colored plum, the C-terminal dimerization domain is colored light green, and disordered flanking and linker regions are colored pale yellow. The second Tr1 chain is colored grey. **(c)** Dimeric H-T-H domain of Tr1 (colored magenta) superimposed onto the crystal structure of HipB bound to its DNA operator from *Shewanella oneidensis*^76^ (colored light blue; PDB: 4PU4). **(d)** Dimeric C-terminal domain of Tr1 (colored green) superimposed onto the crystal structure of EspR from *M. tuberculosis*^77^ (colored brown; PDB: 4NDW)

Next, we used the Dali server^46^ to search the Protein Data Bank (PDB) for structures similar to Tr1, but which have been the subject of structure-function analyses. Amongst the results of the Dali search, nine of the ten structures most similar to Tr1 were DNA-binding proteins with similarity to the N-terminal domain of Tr1 and which function as dimers. Comparing the N-terminal domain of Tr1 with these structures reveal that Tr1 is likely to be a five-helix bundle which contains a DNA-binding helix-turn-helix motif (H-T-H), similar to cI/Cro-like transcription factors (**Fig 3a).**

Prompted by this observation, we next used ColabFold Multimer to generate a model of dimeric Tr1 protein. In the resulting model, the N-terminal domains of Tr1 are closely associated with one another, forming a symmetrical dimer (**Fig 3b).** The C-terminal domains of the two Tr1 monomers, which each resemble two antiparallel helices separated by a turn, interdigitate with one another forming an additional point for dimerization between the two protein chains (**Fig 3b)**. Whilst the global confidence score for the dimeric model of Tr1 was higher than that of the monomeric model (dimer pTM 0.557), the confidence score for the interface of the dimeric Tr1 model was low (ipTM=0.545). We considered that the large proportion of disordered protein in Tr1 could skew the interface confidence scores in our predictions, so we tried to predict dimeric models for Tr1 residues corresponding to the N- and C-terminal domains only. Interface confidence scores for dimeric Tr1 N- or C-terminal domains in isolation were high enough to suggest plausible interactions (piTM=0.763 and 0.69, respectively).

Structures of dimeric H-T-H proteins identified by our Dali search were superimposed onto the Tr1 dimer model to further assess this predicted structure. The N-terminal domain of Tr1 dimerizes in manner typical of other dimeric H-T-H proteins, including those bound to DNA (**Fig 3c**). Frequently, H-T-H proteins that bind DNA as symmetrical homodimers bind to palindromic DNA sequences with each protein binding to a half-site of the palindrome. From these structural comparisons, we predict this is likely to also be the case for Tr1. Fewer structural homologues of the C-terminal domain of Tr1 were identified but we noted similarity between the Tr1 dimer model and the EspR transcription factor of *Mycobacterium tuberculosis* (**Fig 3d)**. EspR dimerizes via a domain similar to the Tr1 C-terminal domain and this is required for EspR to bind DNA with high-affinity^47^.

Next, we used gel filtration chromatography to assess whether purified recombinant Tr1 protein could form dimers or other multimers in solution. Recombinant Tr1 protein was purified and affinity tags removed prior to running on a gel filtration column. We reasoned that removing any additional protein sequences from Tr1 would yield protein most similar to that found in its native environment. Since the expected molecular weight of monomeric recombinant Tr1 is 21.3 kDa, we also ran purified MBP and GFP, predominantly monomeric proteins with respective molecular weights of 43.8 kDa and 28.5 kDa, as controls. Both MBP and GFP eluted from the gel filtration column as monodisperse peaks whereas Tr1 eluted as two overlapping peaks, indicating that Tr1 exists in multiple forms in solution (**Fig 4a).** SDS-PAGE analysis of fractions collected from these experiments confirmed that the peaks contained the expected proteins (**Fig 4b).** Apparent molecular weights for the peaks were calculated with a standard curve based on their elution volumes. The calculated apparent molecular weights for GFP and MBP were 31.6 and 45.7 kDa, respectively, roughly consistent with their expected weights as monomers. The second peak of Tr1 eluted at an apparent molecular weight of 79.4 kDa, suggesting that Tr1 can form tetramers in solution. The first peak of Tr1 eluted at a volume outside of our standard curve and potentially contained aggregated proteins. We generated a predicted tetrameric model of full-length Tr1, but global confidence scores were noticeably lower than those for other Tr1 models (pTM 0.414), so do not present this data here. Further biochemical data are required to guide additional predictive modelling of Tr1 multimers.

**Figure 4.**
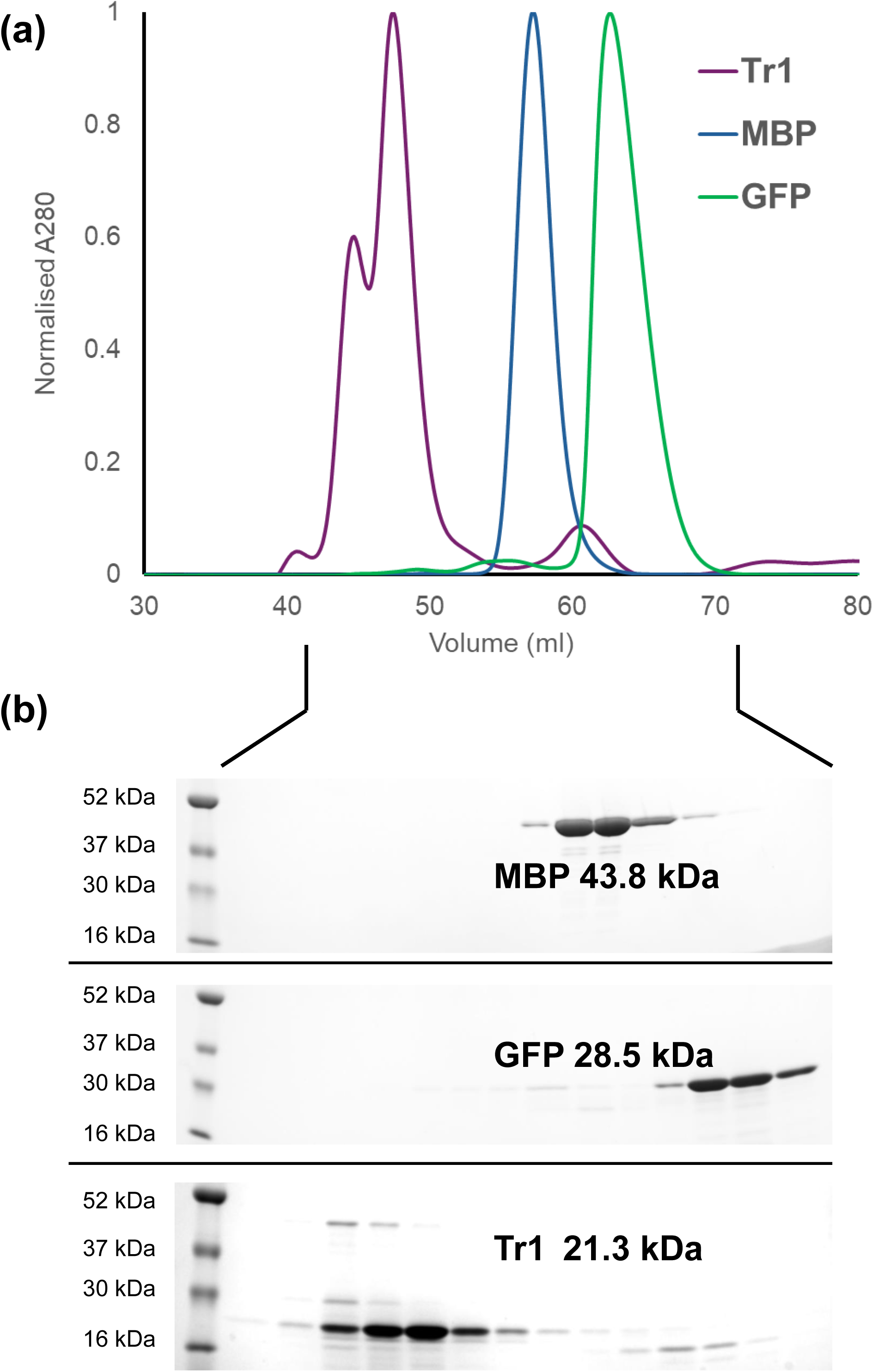
Tr1 forms multimers in solution. **(a)** Chromatograms of Tr1, MBP, and GFP separated by gel filtration. The Tr1 protein lacks tryptophan residues and absorbs weakly at 280 nm so the peaks are shown as normalized absorbance. **(b)** SDS-PAGE gel of fractions collected from equivalent volumes in gel filtration experiments. The left-hand lane of each gel contains molecular weight markers with the known weights labelled.

Taken together, we predict that Tr1 is comprised of H-T-H and dimerization/multimerization domains joined and flanked by regions of disorder. Comparisons indicate that Tr1 might have similarity to cI/Cro transcription factors and EspR from *M. tuberculosis.* Gel filtration experiments demonstrate that Tr1 forms multimers in solution, supporting these predictions, however, a high confidence predicted model of a Tr1 tetramer could not be generated

### Tr1 binds the promotors of neighboring genes omp1X and omp1N

The *tr1* gene was first noticed in studies examining expression of downstream genes *omp1X*, *omp1N*, and the *msp2/p44* expression site^43,44^ (**Fig 5a**). We tested Tr1 binding upstream of *tr1*, *omp1X*, *omp1N*, and the *msp2/p44* expression locus by electrophoretic mobility shift assays (EMSA) using DNA probes for sequences preceding each gene (Fig 5a). EMSA shifts indicated Tr1 complexed with promotors p-*tr1* (**Fig 5b**), p-*omp1X* (**Fig 5c**) and p-*omp1N* (**Fig 5d**). At higher Tr1 concentrations p-*tr1* and p-*omp1N* also displayed secondary complexes suggesting multiple Tr1 binding sites or higher order Tr1 oligomer complexes (**Fig 5b,d**). DNA sequence upstream of the *msp2* expression locus did not shift at any concentration of the Tr1 protein, indicating Tr1 does not individually regulate *msp2* (**Fig 5e**).

**Figure 5.**
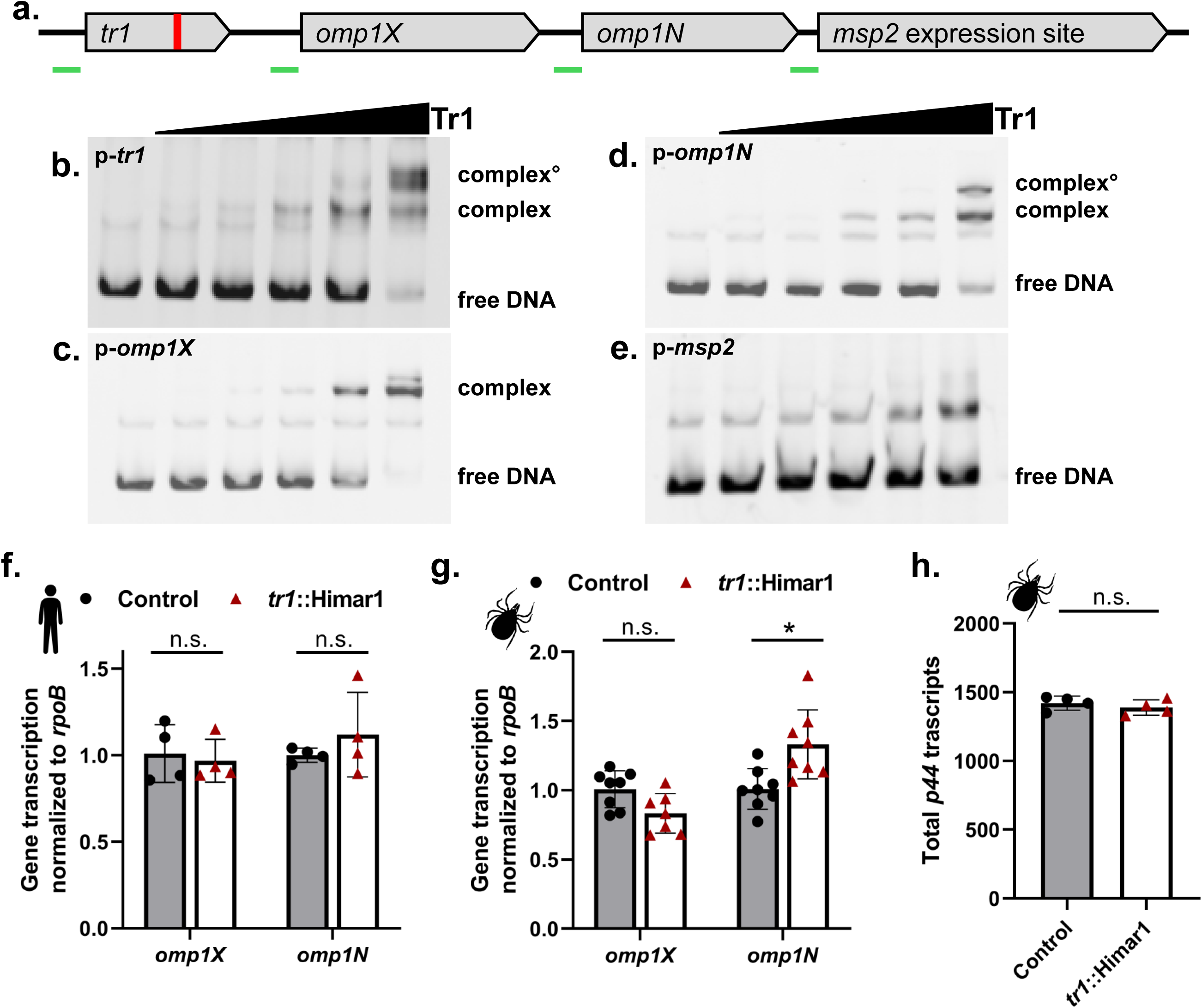
Tr1 binds promotors of neighboring *omp1X* and *omp1N.* (**a**) Diagram of *tr1* gene with downstream neighboring outer membrane protein genes *omp1X*, *omp1N*, and the *msp2/p44* expression site. (**b through e**) EMSA shifts with increasing rTr1 (0, 0.0625, 0.125, 0.25, 0.5, and 1μM ) with DNA probes of (**b**) *tr1*, (**c**) *omp1X*, (**d**) *omp1N*, and (**e**) *msp2/p44* promotor sequences. (**f and g**) Transcription of *omp1X* and *omp1R* from control or *tr1*::Himar1 *A. phagocytophilum* mutant strains at 24 hours post infection in (**f**) human HL60 or (**g**) tick ISE6 cells. Transcripts measured by qRT-PCR and normalized to *rpoB* via ΔΔCt. Data displayed as mean with ±SD of four replicate infections measured with two technical replicates each. \**P* < 0.05 (Mann-Whitney *t*-test). (**h**) *msp2/p44* transcripts totals across all gene variants sequenced by RNAseq from *tr1*::Himar1 or control *A. phagocytophilum* infected ISE6 tick cells. Bars are mean ±SD of four independent infections, each shown as points. n.s. > 0.05 (Mann-Whitney *t*-test).

To ask how loss of *tr1* affected these genes’ transcription, RNA was collected from control and *tr1*::Himar1 *A. phagocytophilum* infected ISE6 cells. Surprisingly, qPCR measuring *omp1X* and *omp1N* transcription from *tr1*::Himar1 and control *A. phagocytophilum* strains (**Fig 5f,g**) found that only *omp1N* displayed a small but significant expression difference during tick cell infection (**Fig 5g**), suggesting Tr1 binding alone does fully account for the behavior of these genes.

### tr1 is required for A. phagocytophilum transcriptional shift during tick cell infection

Tr1’s predicted role as a transcriptional regulator^43,48^ and the inability of the *tr1*::Himar1 to survive in tick cells led us to ask how the loss of *tr1* affects the transcriptome during tick cell infection using RNA sequencing (RNAseq). ISE6 tick cells were infected with *A. phagocytophilum tr1::Himar1* or the control strain. Our previous qPCR for *A. phagocytophilum* gDNA from infected ISE6 cells found that the *tr1*::Himar1 strain persists up to day 3 before declining relative to the control strain (**Fig 1c**). Since measuring genomic DNA (gDNA) may also reflect dead bacteria, we measured *A. phagocytophilum 16s* RNA relative to *I. scapularis actin* transcripts as an indicator of bacterial survival at 12, 24, and 48 hours post infection (hpi). At 24 hours *tr1*::Himar1 survival was 63% of the control strain but fell to only 10% by 48 hpi (**Fig S2**). To capture transcriptional differences when *tr1*::Himar1 bacteria remained viable we performed RNAseq 24 hpi.

Genes downstream of *tr1*, *omp1X*, *omp1N*, and *msp2*^43^ (**Fig 5a**) were all measured in the RNAseq data. Similar to the qRT-PCR measurements (**Fig 5g**), *omp1X* did not significantly differ between *tr1*::Himar1 and the control strain. *omp1N* had a small but significant increase in expression relative to the control (**Table S1**). Measuring *msp2* transcription is complicated by the presence of multiple *msp2* pseudogenes, which *A. phagocytophilum* uses to evade the mammalian antibody response by antigenic variation. For individual *msp2* variants to be transcribed, the pseudogenes recombine into the expression site adjacent to *omp1N*^43,44,49–52^ (Fig 5a). In our RNAseq data, 97 *msp2* variants were detected, with 15 having significant transcription differences between *tr1*::Himar1 and the control strain (**Table S2**). To quantify total *msp2* transcription from the expression site, we summed all *msp2* transcripts from the RNAseq data. This found no difference in overall *msp2* transcription between the *tr1*::Himar1 and control strain (**Fig 5h**). Expression differences among individual *msp2* variants likely reflects *msp2* diversity bottle neck during Himar1 library construction and/or *msp2* recombination since the *tr1*::Himar1 and control strains were purified from the larger collection. These finding again suggest the importance of *tr1* extends beyond its immediate genetic neighborhood.

Beyond *tr1*’s immediate neighbors, RNAseq identified 177 genes as differentially expressed between *tr1*::Himar1 and the control *A. phagocytophilum* strain (p-adj<0.05) during tick cell infection (**Table S1**). Aside from *msp2* variants, 22 genes were >2 fold upregulated in the *tr1*::Himar1, although none have known host or vector-specific expression^32,33^ and are largely ribosomal or other core bacterial genes^34^. Conversely, 14 genes had >2 fold reduced transcription from the *tr1*::Himar1 mutant (**Table 1**). Seven of the of the these, (HGE1_03907/APH_0916, *ateA*, *tr1*, HGE1_01872/APH_0406, *msp4*, HGE1_04767/APH_1111, HGE1_03162/APH_0720) have known tick-specific expression patterns^10,32,33^, which indicates that *tr1* is needed to upregulate *A. phagocytophilum* genes specific for the arthropod vector.

**Table 1.**
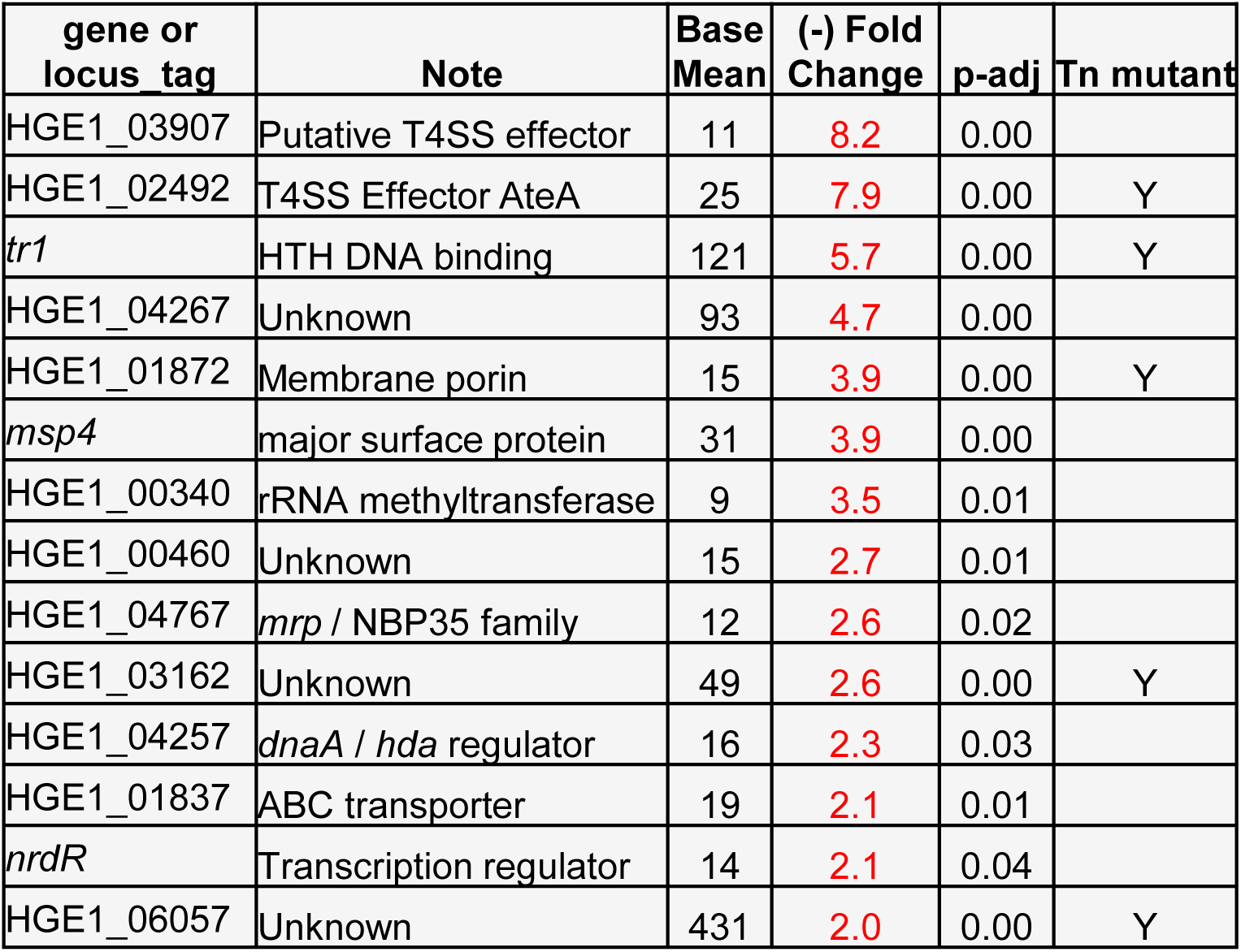
Genes with ≥ 2 fold reduced expression in *tr1*::Himar1 mutant during tick cell infection.

### Tr1 binds promotors of genes of tick-specific A. phagocytophilum genes

We next asked if Tr1 binds promotors of any tick-specific genes impacted by the *tr1*::Himar1 mutation. EMSA were performed with increasing Tr1 protein against promotor sequences preceding; *ateA*, HGE1_06057/APH_1380, HGE1_03162/APH_0720, HGE1_03907/APH_0916, HGE1_01872/APH_0406, and *msp4* (**Fig 6**). Tr1 shifted probes for *ateA* (**Fig 6a**), HGE1_06057 (**Fig 6b**), and *msp4* (**Fig 6c**) at all concentrations, with the majority of the probe band shifted at the highest. HGE1_03162 (**Fig 6d**), HGE1_03907/APH_0916 (**Fig 6e**), and HGE1_01872/APH_0406 (**Fig 6f**) probes also shifted at Tr1 concentrations ≥ 0.125 µM. Similar to EMSAs with probes for *tr1*, *omp1X* and *omp1N,* the higher Tr1 concentrations produced secondary shift bands, suggesting higher order complexes. These findings demonstrate direct Tr1 interaction with promoters for tick-specific *A. phagocytophilum* genes. These EMSA results and differential expression of these genes by the *tr1*::Himar1 mutant strain indicates direct regulation by Tr1.

**Figure 6.**
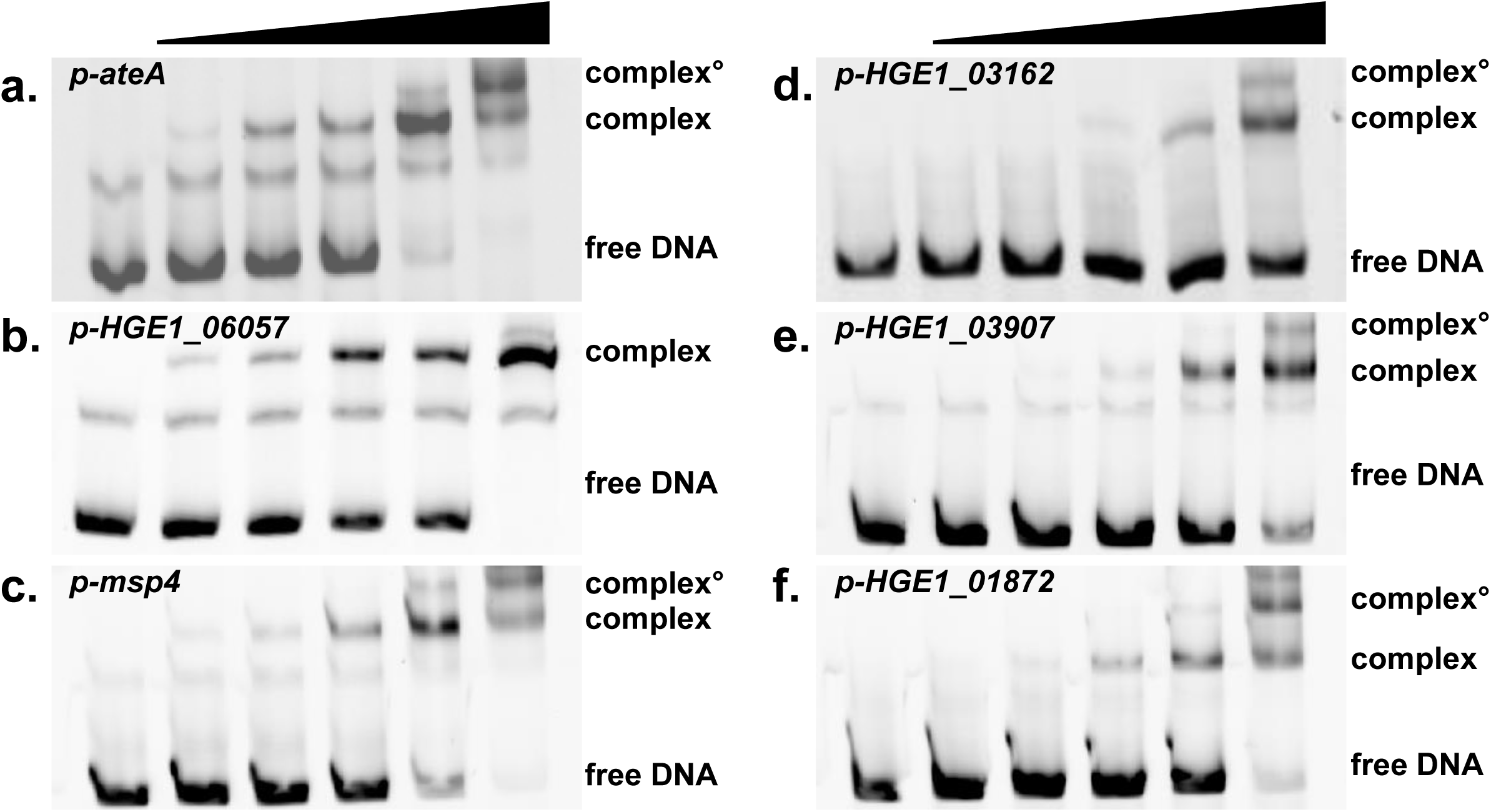
Tr1 binds the promotors of tick-specific *A. phagocytophilum* genes. EMSA shifts with increasing rTr1 (0, 0.0625, 0.125, 0.25, 0.5, and 1μM) tested against promotor sequences of (**a**) *ateA*, (**b**) *HGE1_06057*, (**c**) *HGE1_01872*, (**d**) *HGE1_03162*, (**e**) *HGE1_03907*, and (**f**) *mps4*.

### Tr1 regulated genes are important for A. phagocytophilum survival in tick cells

Inability of the *tr1*::Himar1 to survive in tick cells and the reduced expression of known tick-specific *A. phagocytophilum* genes led us to ask if these genes are important for tick cell infection. We previously showed *ateA*::Himar1 was similarly defective for survival in tick cells and acquisition by ticks^10^. From the *A. phagocytophilum* Himar1 mutant library, we identified and isolated three additional mutants disrupted in genes putatively regulated by Tr1: *HGE1_01872*, *HGE1_03162*, and *HGE1_06057*. Growth of all mutant strains in human HL60 cell culture was comparable to the control strain (Fig 7a). Accordingly, we infected ISE6 tick cells with *A. phagocytophilum* strains *tr1*::Himar1, *ateA*::Himar1, *HGE1_01872*::Himar1, *HGE1_03162*::Himar1, *HGE1_06057*::Himar1, and intergenic control::Himar1. At 8 days post infection all test strains had significantly reduced *Anaplasma* burden in the tick cells relative to the intergenic Himar1 control, with *tr1::Himar1* being the most severe (**Fig 7b**). From this, we demonstrated three additional tick-specific genes HGE1_01872, HGE1_03162, and HGE1_06057 as critical for infecting ticks. Our results suggest that the tick-specific defect of *tr1*::Himar1 mutant strain stems from its failure to express genes controlled by Tr1 and critical for adaptation to the tick cell environment.

**Figure 7.**
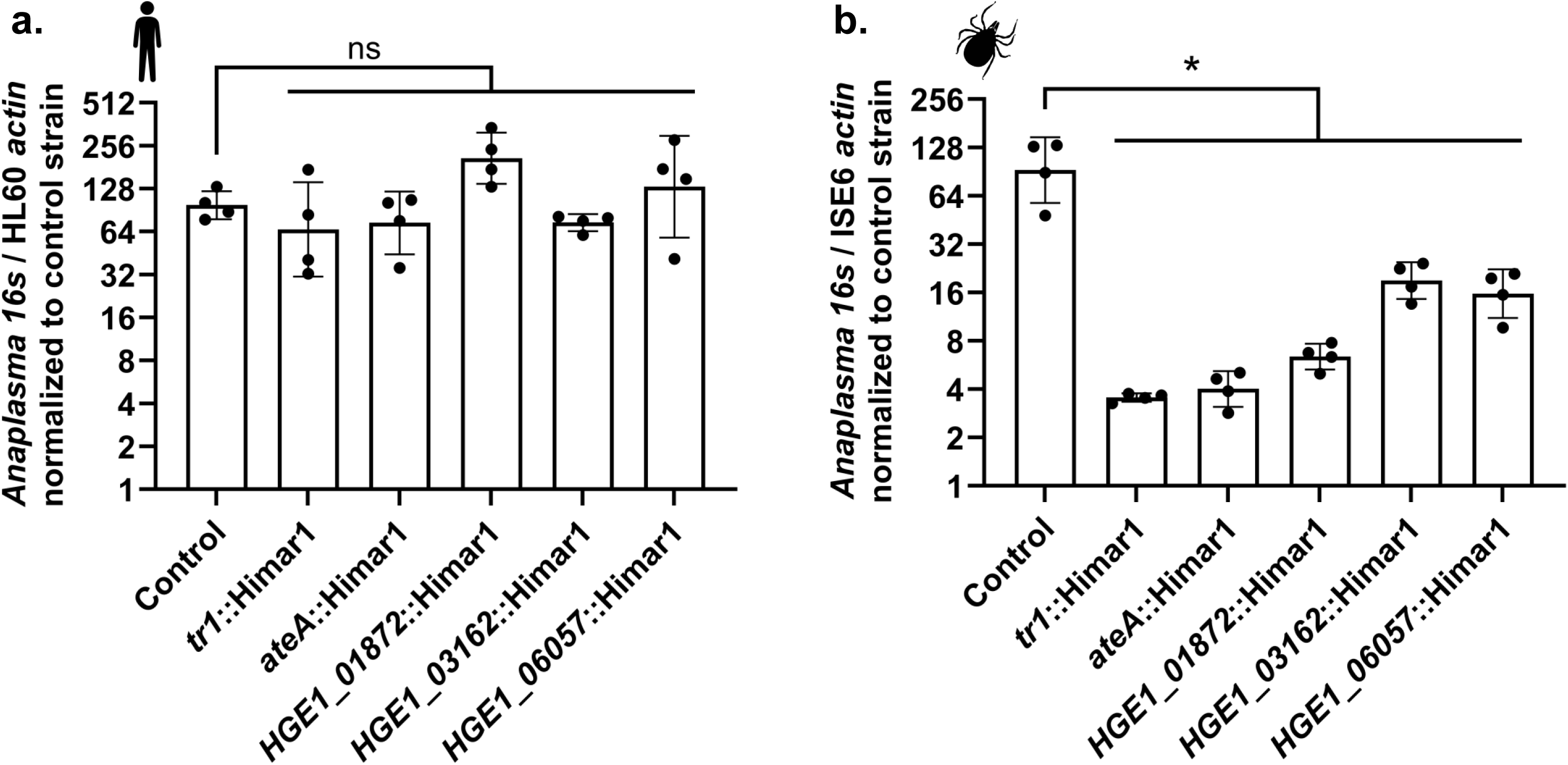
Tr1 impacted genes are necessary for survival during tick cell infection. Survival of indicated Himar1 transposon mutant *A. phagocytophilum* strain relative to intergenic control::Himar1 strain in (**a**) human HL60 (48 hpi) and tick ISE6 cells (8 dpi). Bacterial burden measured by qRT-PCR of *A. phagocytophilum 16s* vs Ixodes actin transcripts. Data displayed as mean ±SD of four replicate infections measured in two technical replicates each. Graph representative of three replicate experiments. * < 0.005 (Mann-Whitney *t*-test).

### tr1 is necessary for tick-specific alterations in the T4SS apparatus

The promotor sequences of *ateA* and HGE1_06057 had high affinity for Tr1 by EMSA (**Fig 6 a,b**). Previously we identified AteA as the first *A. phagocytophilum* T4SS secreted effector specific to tick infection^10^. HGE1_06057 is also predicted as a T4SS substrate by multiple effector prediction algorithms^12,53^. Tr1 binding to promotors of effectors prompted us to ask if Tr1 influences expression of T4SS machinery itself.

An oddity of the *Anaplasma* T4SS is that some components of the apparatus are encoded by multiple paralogs^16,23^. In particular, the needle like pilus of the *A. phagocytophilum* T4SS is encoded for by eight paralogs of the pilin gene *virB2* (**Fig 8a**). Transcriptomics studies noticed different combinations of *virB2* paralogs are upregulated when *A. phagocytophilum* infects mammalian or tick cells^32,33^. Because tiling arrays used in these studies can conflate signal among paralogs with close sequence identity^33^, we first validated *virB2* expression. We designed primers specific to each *virB2* paralog. We grew *A. phagocytophilum* in human HL60s or tick ISE6 cells and measured transcription of each *virB2* paralog by qRT-PCR. As in the transcriptomic studies, mammalian cultured *A. phagocytophilum* only expressed *virB2* paralogs *virB2_1* and *virB2_2* (**Fig 8b**). Conversely, transcription of *virB2_1* and *virB2_2* was reduced during growth in tick cells, while *virB2*_3 → virB2_8 were upregulated (Fig 8b). To examine if the tick-specific expression pattern among the *virB2* paralogs is dependent on *tr1* we reviewed our RNAseq findings for the *tr1*::Himar mutant. Both *virB2_1* and *virB2_2* expression were significantly elevated (1.47 and 1.38 fold) in the *tr1*::Himar1. While not significant due to low number of total reads *virB2_6* and *virB2_7* had reduced expression (0.55 and 0.218 fold) in the *tr1*::Himar1 relative to the control strain (**Table S1**). Together this indicated that the *tr1*::Himar1 strain fails to adjust expression among the *virB2* paralogs during tick cell infection and retains a *virB2* expression pattern similar to mammalian infection.

**Figure 8.**
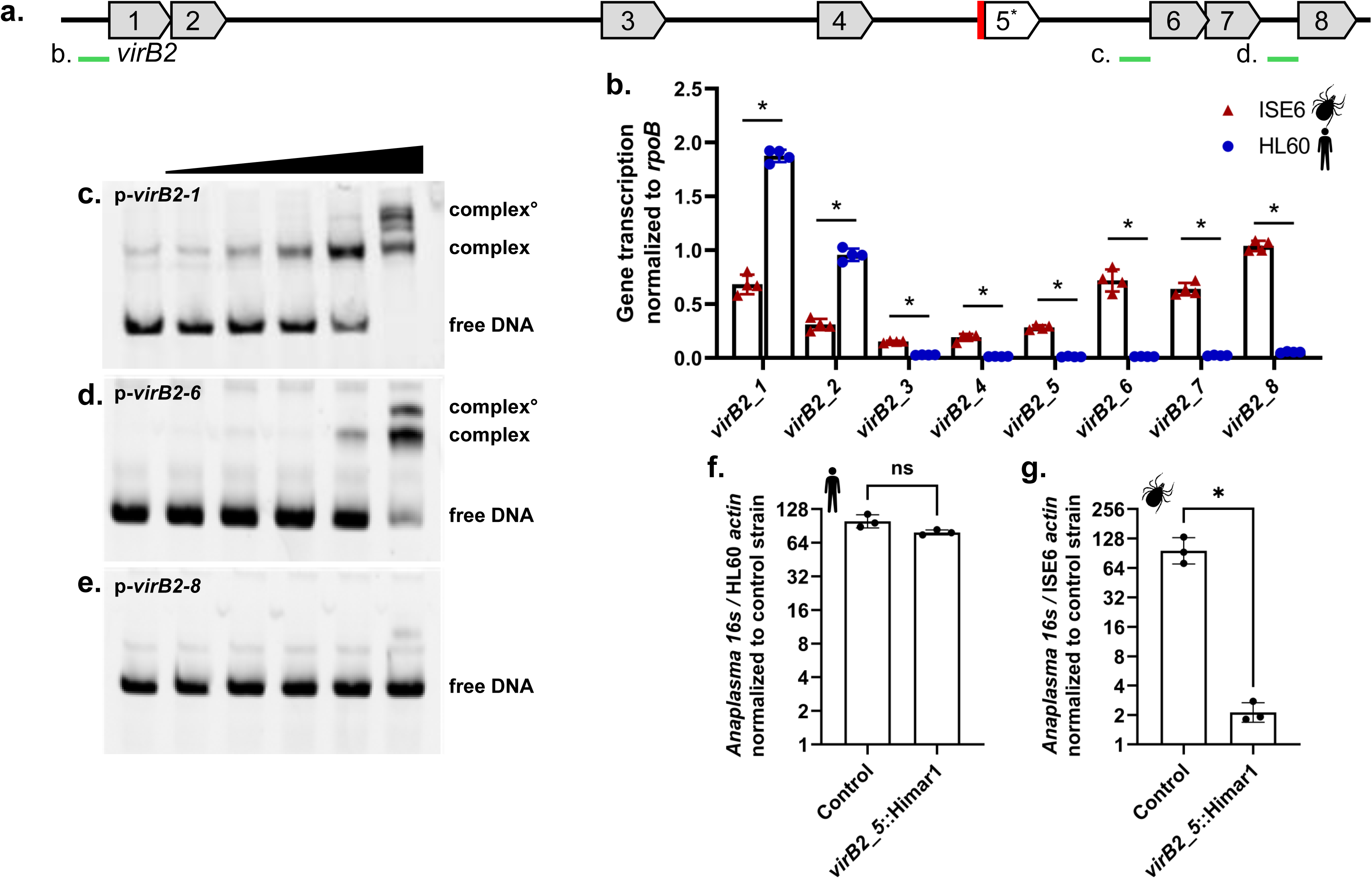
Tr1 is required for tick-specific remodeling of the T4SS pilus. (**a**) Diagram of the *virB2* paralog genes (locus HGE1_04947 – HGE1_04867), the location of the *virB2_5*::Himar1 insertion mutation (red), and potential promotor regions subject to EMSA (green). (**b**) Transcription of the *virB2* genes during wild-type *A. phagocytophilum* infection of mammalian HL60 and tick ISE6 cells. Data shown as mean ±SD of four biological replicates. (**c,d,e**) EMSA shifts with increasing rTr1 (0, 0.0625, 0.125, 0.25, 0.5, and 1μM ) with putative promotor sequences upstream of (**c**) *virB2_1*, (**d**) *virB2_6*, and (**e**) *virB2_8*. (**f and g**) Growth of *A. phagocytophilum virB2_5*::Himar1 or control strain in cell culture infections of (**f**) human HL60 cells and (**g**) tick ISE6 cells. *A. phagocytophilum* burden measured by bacterial gDNA relative to eukaryotic host gDNA via qPCR. Data displayed as mean with ±SD of three biological replicates with two technical replicates each. Data are representative of three experimental replicates. \**P* < 0.05 (Mann-Whitney *t*-test).

### Tr1 binds promotors of T4SS virB2 paralogs differentially expressed in tick infection

We asked if Tr1 can directly bind promotor regions among the differentially expressed *virB2* paralogs. DNA probes designed upstream of *virB2_1*, *virB2_6,* and *virB2_8* (Fig 8a) were tested with increasing concentrations of Tr1 protein by EMSA. Tr1 shifted the promotors of both *virB2_1/2* (**Fig 8c**) and *virB2_6/7* (**Fig 8d**), with *virB2_1/2* shifting completely at the highest concentration. As with other targets, possible secondary complexes were also visible, suggesting possible higher order interactions. Minimal shifting was seen with probes designed upstream of *virB2_8* (**Fig 8e**) suggesting either independent regulation, or co-regulation with the upstream genes *virB_6/7*. Our findings indicate that Tr1 directly participates in regulation of tick-specific expression patterns of T4SS components.

### Tick-specific virB2 paralogs are necessary for A. phagocytophilum survival in tick cells

As with the other Tr1 regulated targets, we asked if the tick-specific *virB2* paralogs are necessary for *A. phagocytophilum* survival in tick cells. From the transposon mutant library, we obtained a mutant strain disrupted in *virB2_5* (*virB2_5*::Himar1) (**Fig 8a**). In the *A. phagocytophilum* HGE1 strain genome (APHH00000000) *virB2_5* is not annotated and therefore was not initially mapped by RNAseq. However, we do detect transcription from the *virB2_5* locus during tick cell infection (**Fig 8b**). The *virB2_5*::Himar1 mutant grew as well as the control strain in human HL60 cells (**Fig 8f**). *A. phagocytophilum virB2_5*::Himar1 and the intergenic control::Himar1 strain were collected from HL60 cell culture and used to infect ISE6 tick cells. Eight days post infection *Anaplasma* burden of *virB2_5*::Himar1 was greatly reduced relative to the control::Himar1 strain in the tick cells (**Fig 8e**). This finding indicates that tick-specific *virB2* paralogs are important for *A. phagocytophilum* adaptation to the tick.

### Tr1 homologs are found throughout the Rickettsiales

Having established a role for Tr1 in the survival of *A. phagocytophilum* within the tick host, we queried whether Tr1 homologs are present in other Rickettsiales^43^, since many of these bacteria also have life cycles that involve cycling between vertebrate and arthropod hosts. Using sequence-based searches within the Rickettsiales, we identified candidate homologs of Tr1 in diverse Anaplasmataceae genomes. The *tr1* genes of *Anaplasma*, *Ehrlichia*, and *Neoehrlichia* share a high degree of genomic synteny and are invariably co-localized with *ndk* (encoding nucleoside-diphosphate kinase) and one or more OMP-encoding genes. In *Wolbachia* and *“Candidatus* Mesenet*“* the *tr1* locus is more variable with *ndk* being lost and OMP-encoding genes being co-localized at the *tr1* locus in some, but not all, species.

We used Foldseek to extend our search for more distant Tr1 homologs, filtering our results for Rickettsiales bacteria. Using this approach, we identified Tr1 proteins in *Rickettsia* and *“Candidatus* Tisiphia*“* species. Whilst the *tr1* locus in *Rickettsia* species is not syntenic with those from the Anaplasmataceae, the *Rickettsia tr1* gene is most often co-localized with *sca4.* Intriguingly, *sca4* has been identified as having a role in fitness of *Rickettsia parkeri* within the tick host^54^. Yet more distant Tr1 homologs were also identified in *Midichloria* and *Orientia*, although in the latter the C-terminal domain is truncated or lost. Candidate *tr1* homologues are co-localized with genes that encode proteins with unknown functions in many Rickettsiales.

These results indicate that Tr1 represents a family of transcription factors found broadly across the Rickettsiales, suggesting that some of our findings could be extrapolated to other bacteria of this Order.

## DISCUSSION

As a vector-borne pathogen, survival in both the tick and the mammal is essential to complete the transmission cycle of *A. phagocytophilum* and be maintained in the environment. We know the bacteria undergoes extensive transcriptional changes between the two environments^32,33^ and have identified individual adaptations that are critical for either mammalian or tick infection^34–37^. Understanding how rickettsial pathogens switch their host tropism to the arthropod, could inform interventions to disrupt the transmission cycle. In this work we identified Tr1 as a critical switch regulating *A. phagocytophilum* adaptation to the tick and identified Tr1-controlled genes necessary for survival in the arthropod. Collectively our findings uncover a central node in a network of changes necessary for rickettsial bacteria to complete their vector-borne lifecycle.

Due to the obligate intracellular lifecycle and lack of compatible extrachromosomal plasmids, genetic manipulation of *A. phagocytophilum* remains challenging. The Himar1 transposon library has proven to be an invaluable resource, allowing the first phenotypic testing of gene disruptions in *Anaplasma*^34^. The library was generated in HL60 cell culture, which prohibits mutation in genes essential for survival in mammalian cells. However, *A. phagocytophilum* genes specific to tick infection were readily mutated^34,35^. Such mutants have been leveraged to uncover tick-specific roles of the T4SS effector AteA^10^, duplicate T4SS component VirB6-4^36^, and surface protein modifying O-methyltransferase^37^. In our study we extensively relied on mutants from this library to uncover tick-specific contributions of *tr1* and five of the genes it impacts. Until more targeted genetic tools are available in this system, the *A. phagocytophilum* Himar1 transposon library remains a critical resource.

Initial interest in the *tr1* gene stemmed from its location upstream of outer membrane proteins genes *omp1X*, *omp1N*, and *msp2/p44* expression site^43,44^. ApxR was identified as the first *Anaplasma* DNA-binding transcriptional regulator, and was later shown to bind its own promotor and upstream region of the *p44/msp2* expression site. Similarly, the ApxR homolog EcxR in the related rickettsial pathogen *Ehrlichia chaffeensis*, binds promotors of *tr1* and downstream outer membrane protein genes^55,56^. *E. chaffeensis* Tr1 also binds upstream membrane protein genes *p28* and *omp-1B*^55^. We found *A. phagocytophilum* Tr1 binds DNA sequences upstream of *omp1X* and *omp1N*. However, only expression of *omp1N* differed between the *tr1*::Himar1 mutant and control strain, indicating *tr1* is not the only factor controlling these genes. Though the number, identity, and arrangement of adjacent membrane protein genes downstream of *tr1* in *Anaplasma* and *Ehrlichia* differ greatly^44,55^, common models may emerge showing how Tr1 and ApxR/EcxR participate in regulation of the *tr1* adjacent surface protein genes.

Looking past its immediate chromosomal neighbors multiple tick-specific membrane protein genes had reduced expression in the Himar1::*tr1* and were directly bound by Tr1. These include; *HGE1_03162* (*Aph_0720*), *HGE1_01872* (*Aph_0406*), *msp4*, and *HGE1_03907* (*Aph_0916*). Transcription of all four is highly specific to *A. phagocytophilum* infection in tick cells^32,33^. Reflecting the expression pattern, we found *HGE1_03162* and *HGE1_01872* are necessary for tick cell infection. Adjacent to *HGE1_01872, A. phagocytophilum* encodes two *HGE1_01872* paralogs *asp62* (*HGE1_01862, Aph_0404*) and *asp55* (*HGE1_01867, Aph_0505*). Antibodies against Asp62 and Asp55 were found to limit *A. phagocytophilum* infection of human HL60 cells^57^. While Asp62 and Asp55 are expressed during both mammalian and tick cell infection, HGE1_01872 is specific to the arthropod vector^32,33^. Both HGE1_01872 and Msp4 are substrates of an O-methyltransferase required for tick infection and promote binding to *Ixodes* cells^37,58,59^. Similar to HGE1_01872, HGE1_03907 (Aph_0916) is adjacent to the invasion protein gene *aipA*, which is necessary for infection of mammalian cells^60^, suggesting HGE1_03907 represents a arthropod-specific AipA alternative. These findings implicate Tr1 as the switch regulator governing this remodeling of the *A. phagocytophilum*’s surface proteome necessary for tick infection.

Beyond surface protein genes, Tr1 is playing a broader role controlling genes expressed during infection of the tick. Our findings identify Tr1 as a critical regulator of tick-specific effector AteA and specialization of the T4SS VirB2 pilus necessary for adaptation to the tick. The full repertoire of effectors secreted by *A. phagocytophilum* is unknown, but effector predicting algorithms propose as many as 49^12,53^. So far, few have been experimentally examined^17,21,22,24,61–63^, with only AteA investigated in context of the arthropod vector^10^. The *E. chaffeensis* ApxR homolog, EcxR, was also shown to regulate components of the T4SS apparatus during mammalian cell infection^64^. However, EcxR has yet to be examined during tick infection, and binding to effector to *virB2* pilin genes is untested. Uncovering the full regulome of Tr1 and how it works with other regulators to manage adaptation between the mammalian and tick environments will likely require larger unbiased binding and regulation studies.

How does Tr1 recognize its DNA targets and regulate gene expression? Bioinformatic analysis indicates that the Tr1 protein has similarity to H-T-H transcription factors and, inferred from this and our protein modelling, is likely to dimerize and bind palindromic DNA motifs to exert influence on transcription on downstream genes. Consistent with this, we also identified a putative dimerization domain in the C-terminus of Tr1 and we observed that recombinant Tr1 protein forms multimers in solution, probably tetramers and possibly other higher order structures. Other H-T-H proteins such as the λ repressor, share this ability to form multimers of dimers, enabling them to bind to multiple DNA sites co-operatively. Indeed, multiple shifts of Tr1-DNA complexes in our EMSAs, supporting that binding of Tr1 might be co-operative at some of its target sites. Understanding if or how the multimerization of Tr1 is regulated during infection, the interplay between Tr1 and other transcription factors at promoter regions, and the identification of a consensus DNA target site and regulon will be crucial for determining how Tr1 mediates *A. phagocytophilum* adaptation to the arthropod host.

We identified candidate Tr1 homologs throughout the Rickettsiales. In both *Anaplasma* and some *Rickettsia,* the Tr1 gene is co-localized with genes that are either differentially expressed between the tick and vertebrate host or have been implicated in survival in tick cells. This functional conservation of the *tr1* loci between these two species makes it interesting to consider that Tr1 might have a role in transcriptional remodeling that underpins the cycling between vertebrate and arthropod hosts in diverse Rickettsiales. However, this leaves the question of what the role of Tr1 in Rickettsiales that survive exclusively within arthropods, such as *Wolbachia,* is. Furthermore, in many Rickettsiales, the *tr1* gene is co-localized with genes of no known function. Whilst genomic synteny suggests that Tr1 fulfills similar roles in *Anaplasma, Ehrlichia* and *Neoehrlichia,* further study is required to determine how Tr1 functions across diverse Rickettsiales.

Altogether, our work identifies Tr1 as an essential master regulator that remodels the *A. phagocytophilum* transcriptome to infect the tick vector. Further, we identified multiple Tr1 regulated genes that contribute to tick cell infection. With most studies focusing on how *A. phagocytophilum* infects mammals, this work provides insights into an understudied aspect of pathogen biology, uncovering wide-reaching transcriptional events that underpin adaption to the arthropod vector. Unable to be transmitted vertically to host progeny, *A. phagocytophilum* must cycle between the host and tick vector to be maintained in the environment. Tr1 and the genes under its control are an essential part of this system. Uncovering how *A. phagocytophilum* and other rickettsial bacteria achieve this is key to developing interventions to interrupt the transmission cycle.

## MATERIALS AND METHODS

### Bacterial and eukaryotic cell culture

*Escherichia coli* used for plasmid construction, plasmid amplification, and protein expression was cultured with solid and liquid lysogeny broth (LB) medium with the addition of kanamycin or zeocin 25 µg ml^−1^ antibiotics as needed to select for plasmid encoded resistance genes. For production of recombinant Tr1 proteins, *E. coli* strain DE3 (New England Biolabs) was used with LB or Terrific Broth (TB) supplemented with 100 µg ml^−1^ kanamycin.

HL60 human promyelocytic cells (ATCC; CCL-240) were maintained in Roswell Park Memorial Institute (RPMI) 1640 medium with 10% FBS and 1× Glutamax. HL60 cultures were kept at 37°C with 5% CO_2_ in a humidified incubator. HL60 cell cultures were limited to < 20 passages to prevent phenotypic drift and kept between 5 × 10^4^ and 1 × 10^6^ cell/ml to prevent differentiation.

Wild-type *A. phagocytophilum* strain HGE1 and HGE1 derived Himar1 transposon mutants were grown in HL60 cells as previously described^34^. All Himar1 mutant strains were obtained from the previously established *A. phagocytophilum* mutant collection^34^. Status of *A. phagocytophilum* infections in HL60 cells was monitored by Diff-Quick Romanowsky–Giemsa staining. To generate host-cell-free *A. phagocytophilum,* peak infected HL60 cultures (>95% infected cells) were gently sonicated to disrupt HL60 cell membranes and liberated bacteria were separated from host cell debris by differential centrifugation. *Anaplasma* numbers were estimated as previously described ^65,66^.

*I. scapularis* (Say) embryonic tick cells (ISE6) were maintained in L15C-300 medium with 10% FBS (Sigma; F0926), 10% tryptone phosphate broth (TPB; BD; B260300) and 0.1% lipoprotein cholesterol concentrate (MP Biomedicals; 219147680)^67^, in sealed tissue culture treated flasks, and incubated at 34°C ^68^. Infected ISE6 cell cultures were additionally supplemented with 0.25% NaHCO_3_ and 25 mM HEPES buffer (Sigma), cultured in vented flasks, and kept at 34°C in a humidified chamber with 4% CO_2_.

### A. phagocytophilum mutant growth curves

*A. phagocytophilum* burden in HL60 and ISE6 cells was evaluated as previously described^36,37^. Briefly, HL60 cells were seeded to 24 well plates at 5×10^4^ cells/well and infected at an MOI of 1. Three to four replicate wells were harvested at the time of inoculation and tested time post inoculation. ISE6 cells were seeded to 24 well plates at 3×10^5^ cells/well. After ISE6 cells adhered to the plate for at least 6 hours, they were inoculated at an MOI of 50 with host-cell-free *A. phagocytophilum* purified from HL60 cultures. Bacteria were allowed to infect the tick cells for 4 hours, after which the media was exchanged three times to remove excess extracellular bacteria. Four replicate wells were collected post media exchanged and subsequent time points. Pelleted samples were frozen and later processed for RNA or gDNA using Zymogen Quick-RNA Micro-prep or QIAGEN DNeasy blood and tissue kits. Ratios of bacterial and host cell gDNA copies was measured by qPCR as previously described^10,36,37^ Levels of live *A. phagocytophilum* Himar1 mutant bacteria in each infected ISE6 or HL60 cell sample was measured by qRT-PCR targeting *A. phagocytophilum* 16S rRNA and *I. scapularis* or human *β-actin* transcripts (**Table S3**) and compared relative to the intergenic Himar1 control strain by ΔΔCt^10,65,69^.

### Measuring A. phagocytophilum gene transcription

HL60 and ISE6 cells in 24 well plates were infected with wild-type *A. phagocytophilum* HGE1 as described above. At 24 hours post infections four wells were collected and processed for RNA Zymo Quick-RNA micro-prep kit^®^ (ZymoResearch) according to product protocols for tissue culture samples. cDNA was generated with Verso cDNA Synthesis Kit (ThermoFisher). Transcripts of *A. phagocytophilum* genes of interest were measured by qPCR using gene specific primers (**Table S3**) and SYBR green iTaq universal Supermix (Bio-Rad; 1725125) according to Bio-Rad specified cycle conditions. Genes of interest were measured relative to housekeeping gene *rpoB* and compared between human and tick cell infections by ΔΔCt.

### Animal infection

All mice were purchased as six weeks of age from The Jackson Laboratory. Gender balanced groups of ten C57BL/6 mice were infected as previously described^10^ with either the *tr1*::Himar1 or the intergenic Himar1 mutant *A. phagocytophilum*. The intergenic Himar1 mutant was previously shown to be phenotypically equivalent to wild-type^10,36,37^. Blood was collected seven days post inoculation and *A. phagocytophilum* burden in the blood was measured by qPCR as in prior studies (16S rRNA / mouse *β-actin*^10,65,69^) (Table S3). Larval *I. scapularis* ticks purchased from Oklahoma State University (Stillwater, OK, USA) were kept at 23°C with >95% humidity and a 16/8-h light/dark photoperiods. Mice were selected from the intergenic Himar1 and *tr1*::Himar1 strain infected mice with equivalent *Anaplasma* burden. Each mouse was infested with 200 naïve larval *I. scapularis* and allowed to feed to repletion. Detached replete ticks were individually processed and assayed for *A. phagocytophilum* burden by qRT-PCR measuring 16S rRNA versus *I. scapularis β-actin* transcripts (Table S3), compared by absolute quantification ^10,65,69^. All animal use protocols were approved by the Washington State University Institutional Animal Care and Use Committee (ASAF #6630) and animal housing facilities at Washington State University in Pullman, WA maintains AAALAC-accreditation.

### RNA sequencing

ISE6 cells were seeded to 12 well plates at 5×10^5^ cells/well and allowed to adhere for 6 hours. The intergenic transposon control and Himar1::*tr1* mutant *A. phagocytophilum* strains were prepared as host-cell-free bacteria from >95% infected HL60 cultures immediately before ISE6 infection. Adhered ISE6 cells were infected at an MOI of 100. Twenty-four hours post infection media was exchanged three times to remove excess extracellular bacteria, and four replicate wells were collected and RNA isolated with Zymogen Quick-RNA Mini-prep kits. RNA was submitted to the WSU Genomics Core for total RNA-seq analysis. Sequencing libraries were prepared using Tru-seq Stranded Total RNA kit, with Ribo-Zero Plus and libraries, sequenced using a HiSeq 2500, and results exported as FASTQ files as previously discribed^70^. RNA-seq data were aligned with the *A. phagocytophilum* HGE1 genome (GCF_000478425.1)^71^. Transcript quantification and differential gene expression were analyzed by HTSeq and DESeq2^72^.

### Overexpression and purification of recombinant Tr1 protein

The *tr1* open reading frame, codon optimized for expression in *E. coli,* was synthesized and cloned by Twist Bioscience into a modified pET28a expression vector. The resultant plasmid encoded the Tr1 protein fused at its N-terminus to a hexa-histidine tagged maltose binding protein (MBP) with a tobacco etch virus (TEV) protease recognition site between Tr1 and MBP. The plasmid was transformed into *E. coli* DE3 for overexpression of the recombinant Tr1 protein. Briefly, overnight cultures, grown at 37 °C in selective LB media, were used to inoculate flasks of TB. Over-expression of MBP-Tr1 was induced by the addition of 0.5 mM IPTG once cultures had reached an OD_600_ of 0.5-0.6 and then grown for an additional 18 hours at 25 °C. Cells were harvested by centrifugation and kept at -20 °C for short-term storage.

To purify MBP-Tr1 protein, cell pellets were resuspended in Buffer A (20 mM imidazole, pH 8.0; 20 mM Tris-HCl, pH 8.0; 400 mM NaCl; 2 mM beta-mercaptoethanol) and then lysed by sonication. Lysates were clarified by centrifugation and then loaded onto a HisTrap HP purification column (Cytiva). The column was washed extensively with Buffer A prior to elution of the His-tagged MBP-Tr1 with stepwise (10, 20, 100%) washes with Buffer B (as Buffer A but with 400 mM imidazole). Fractions containing MBP-Tr1 were pooled, concentrated by centrifugal-filtration, then further purified by gel-filtration on a Superdex 200 column (Cytiva) equilibrated in SEC Buffer (20 mM Tris-HCl pH 8.0; 250 mM NaCl; 2 mM beta-mercaptoethanol). Fractions containing MBP-Tr1, which eluted as a single peak from the gel-filtration column, were supplemented with his-tagged TEV protease and incubated at 4 °C for 16 hours to cleave the His-tagged MBP protein from Tr1. The resulting protein preparation was passed over a HisTrap HP column which was then washed step (5, 10, 100%) with Buffer B. Tr1 protein typically passed through the HisTrap HP column or eluted in 5% Buffer B washes and were thus separated from the His-tagged TEV and MBP proteins, which eluted in 100% Buffer B. This purification strategy yielded highly pure Tr1 protein lacking any tags.

### Gel filtration

Purified tagless Tr1 protein, green fluorescent protein (GFP) and maltose binding protein proteins were analyzed by gel filtration using a HiPrep Sephadex 75 16/600 column (Cytiva), using Buffer B. GFP and MBP were expressed using pET vectors (modified pET41c and pET28a, respectively) in *E. coli* DE3 and purified by nickel affinity chromatography or amylose affinity chromatography using standard methods.

### Electrophoretic-Mobility-Shift-Assays

Promoter proximal DNA sequences for use in Electrophoretic-Mobility-Shift-Assays (EMSAs) were amplified by PCR. Primers for this purpose were designed to yield DNA fragments comprising the ATG start codon of a selected gene plus 250 bp of upstream DNA sequence. In each case, the forwards primer was appended with an M13 sequence upstream of the primer sequence (5’- GAG CGG ATA ACA ATT TCA CAC AGG). The resulting DNA fragments were separated by agarose gel electrophoresis and fragments of the anticipated size were excised and purified (QIAquick Gel Extraction Kit). These purified fragments were then used as the templates for a secondary PCR using an M13 sequence primer labelled with FAM at the 5’ end in conjunction with the fragment-specific reverse primer used in the primary PCR. The resulting FAM-labelled PCR products were resolved by agarose gel electrophoresis and purified as above. Sequences of primers used are included in **table S3**.

EMSA assays were performed using a modified protocol described elsewhere^73^. Briefly, for each DNA fragment an EMSA mixture comprising (100 ng FAM-labelled DNA fragment;200 ng of herring sperm DNA; 5% glycerol; 10 mM Trs-HCl, pH 8.0; 25 mM KCl) was prepared and 8 µl of this dispensed into a series of tubes. These were then supplemented with 2 µl of purified Tr1 protein at varying concentrations, or with buffer alone, and then mixed by gentle pipetting. Tr1 protein was used at final concentrations of 0.25, 0.5, 1, 2, and 4 µM for all EMSAs presented here. Resulting mixtures were incubated at 25 °C for 30 minutes before being loaded onto 7.5 % polyacrylamide gels buffered with 0.25x TBE and 5% glycerol. Gels were run for 60 minutes at 100 V and then fluorescent images (green channel) were collected on a ChemiDoc MP Imaging System (Bio-Rad).

### Bioinformatics

Protein structures were analyzed and visualized in Coot^74^ and ChimeraX^75^. Modelling and the generation of predicted structures were done using ColabFold and AlphaFold Server with searches for homologues in the PDB being achieved with the Dali Server^45,46^.

---

**Figure S1.**
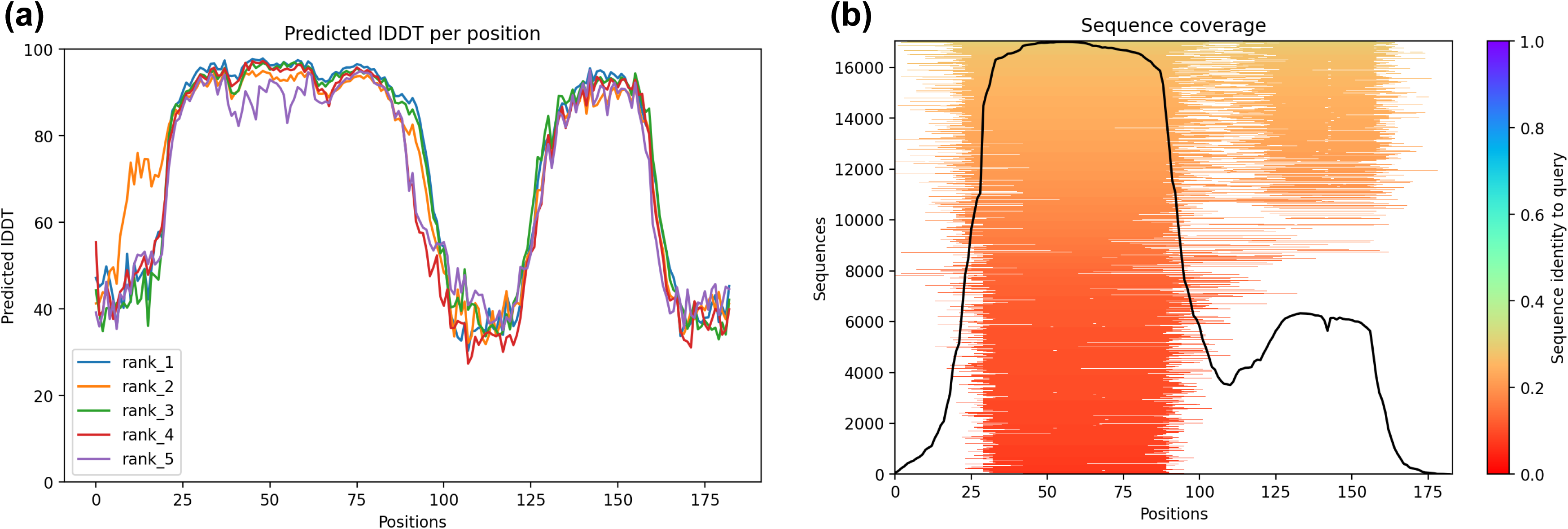
Predicted Local Distance Difference Test (pLDDT) and Multiple Sequence Alignment (MSA) plots from ColabFold of Tr1. (a) pLDDT plot shows that local confidence in the Tr1 model is high in the N- and C-terminal domains, supporting the proposal that Tr1 is comprised of two ordered domains. (b) MSA plot showing that the Tr1 N- and C-terminal domains are evolutionarily conserved.

**Figure S2.**
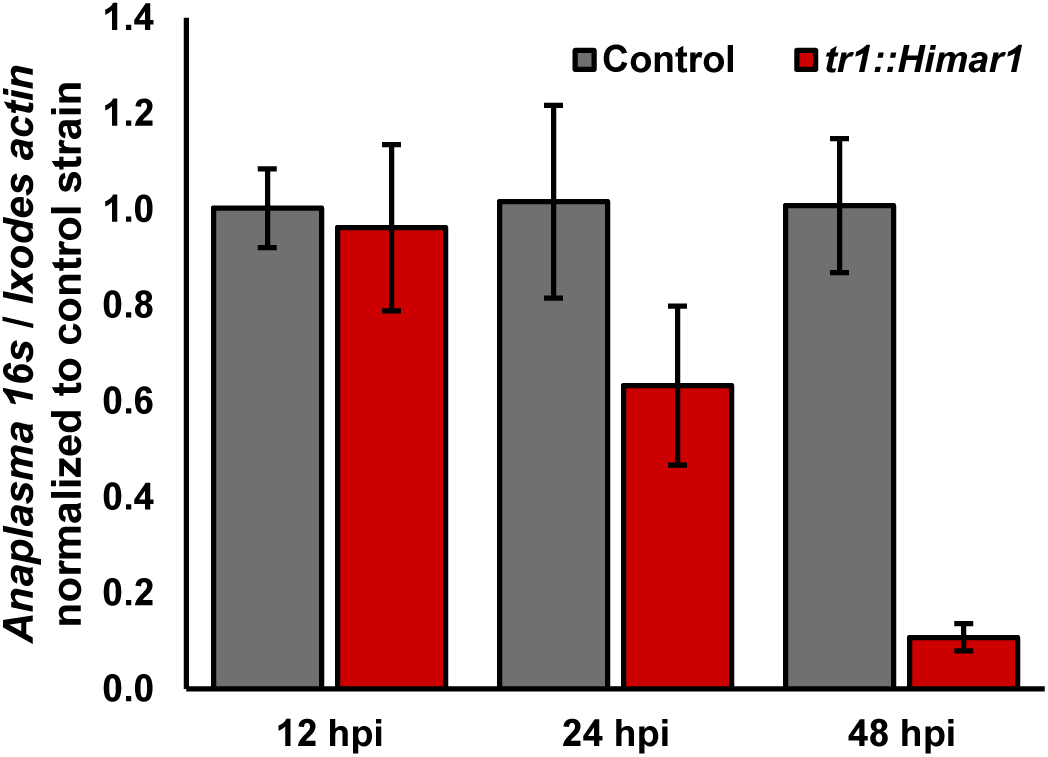
Early survival of *tr1*::Himar1 strain during tick cell infection. *A. phagocytophilum* tr1::Himar1 or control strain in cell culture infections of tick ISE6 cells. *A. phagocytophilum* burden measured by bacterial 16s relative to tick *actin* transcripts via qRT-PCR. Data displayed as mean with ±SD of three biological replicates. Data are representative of three experimental replicates. *P < 0.05 (Mann-Whitney t-test).

## ACKNOWLEDGEMENTS

We are grateful to Ulrike Munderloh, Lisa Price and Nicole Burkhardt at the University of Minnesota (Saint Paul, MN) and Kelly Brayton at Washington State University (Pullman, WA) for providing strains from the Himar1 transposon and methodological guidance. Kelly Brayton for sharing resources. Deirdre Fahy provided administrative support. Washington State University (WSU) Genomics Core performed RNA-sequencing and analyses.

## FUNDING

This work was funded through generous support from; the National Institutes of Health (NIAID grant numbers R21AI178392, R21AI154023, R21AI151412, R61AI179933, R01AI162819), Washington State University Intramural College of Veterinary Medicine grants program funded by the Marvel Shields Autzen Fund and Stanley L. Adler Research Fund, and Washington State University, College of Veterinary Medicine. The Cadby laboratory was also funded by Royal Society Research Grant RGS\R1\231117 and Academy for Medical Sciences Springboard Award SBF009\1204.

## AUTHOR CONTRIBUTIONS

**EricaRose Warwick:** Methodology, Investigation, Visualization, Reviewing & Editing, R**achel Burt:** Methodology, Investigation, Visualization, **Jeffrey T. Badigian:** Methodology, Investigation, **Daniel Howell:** Methodology, Investigation, **Kyle T. Swallow:** Methodology, Investigation, **Chloe Leach:** Methodology, **Azeza M. Falghoush:** Methodology, **Dana K. Shaw:** Methodology, Investigation, Review & Editing. **Ian T. Cadby:** Conceptualization, Supervision, Methodology, Investigation, Writing, Visualization, Funding acquisition. **Jason M. Park:** Conceptualization, Supervision, Methodology, Investigation, Writing, Visualization, Funding acquisition.

## Notes

### Competing Interest Statement

The authors have declared no competing interest.

### Summary of Updates

Correction made to figure 8 f and g

## REFERENCES

1. Diop A, Raoult D, Fournier PE. Rickettsial genomics and the paradigm of genome reduction associated with increased virulence. Microbes Infect. 2018 Aug 1;20(7):401–409.

2. Brayton KA, Kappmeyer LS, Herndon DR, Dark MJ, Tibbals DL, Palmer GH, McGuire TC, Knowles DP. Complete genome sequencing of *Anaplasma marginale* reveals that the surface is skewed to two superfamilies of outer membrane proteins. Proc Natl Acad Sci U S A. 2005 Jan 18;102(3):844–849. PMCID: PMC545514

3. Mavromatis K, Doyle CK, Lykidis A, Ivanova N, Francino MP, Chain P, Shin M, Malfatti S, Larimer F, Copeland A, Detter JC, Land M, Richardson PM, Yu XJ, Walker DH, McBride JW, Kyrpides NC. The genome of the obligately intracellular bacterium *Ehrlichia canis* reveals themes of complex membrane structure and immune evasion strategies. J Bacteriol. 2006 June;188(11):4015–4023. PMCID: PMC1482910

4. McLeod MP, Qin X, Karpathy SE, Gioia J, Highlander SK, Fox GE, McNeill TZ, Jiang H, Muzny D, Jacob LS, Hawes AC, Sodergren E, Gill R, Hume J, Morgan M, Fan G, Amin AG, Gibbs RA, Hong C, Yu XJ, Walker DH, Weinstock GM. Complete genome sequence of *Rickettsia typhi* and comparison with sequences of other rickettsiae. J Bacteriol. 2004 Sept;186(17):5842–5855. PMCID: PMC516817

5. Hammac GK, Pierlé SA, Cheng X, Scoles GA, Brayton KA. Global transcriptional analysis reveals surface remodeling of *Anaplasma marginale* in the tick vector. Parasit Vectors. 2014 Apr 21;7:193. PMCID: PMC4022386

6. Rurangirwa FR, Stiller D, Palmer GH. Strain diversity in major surface protein 2 expression during tick transmission of *Anaplasma marginale*. Infect Immun. 2000 May 1;68(5):3023–3027. PMID: 10769008

7. Kahlon A, Ojogun N, Ragland SA, Seidman D, Troese MJ, Ottens AK, Mastronunzio JE, Truchan HK, Walker NJ, Borjesson DL, Fikrig E, Carlyon JA. *Anaplasma phagocytophilum* Asp14 Is an invasin that interacts with mammalian host cells via its C terminus to facilitate infection. Infect Immun. 2013 Jan;81(1):65–79. PMCID: PMC3536139

8. Lind MCH, Naimi WA, Chiarelli TJ, Sparrer T, Ghosh M, Shapiro L, Carlyon JA. *Anaplasma phagocytophilum* invasin AipA interacts with CD13 to elicit Src kinase signaling that promotes infection. mBio. 15(10):e01561–24. PMCID: PMC11481542

9. Gaywee J, Radulovic S, Higgins JA, Azad AF. Transcriptional analysis of *Rickettsia prowazekii* invasion gene homolog (invA) during host cell infection. Infect Immun. 2002 Nov;70(11):6346–6354. PMCID: PMC130406

10. Park JM, Genera BM, Fahy D, Swallow KT, Nelson CM, Oliver JD, Shaw DK, Munderloh UG, Brayton KA. An *Anaplasma phagocytophilum* T4SS effector, AteA, is essential for tick infection. mBio. 2023 Oct 31;14(5):e0171123. PMCID: PMC10653876

11. Beyer A, Truchan H, Levi M, Walker N, Borjesson D, Carlyon J. The *Anaplasma phagocytophilum* effector AmpA hijacks host cell SUMOylation. Cell Microbiol. 2014 Oct 1;17.

12. Esna Ashari Z, Brayton KA, Broschat SL. Prediction of T4SS effector proteins for *Anaplasma phagocytophilum* using OPT4e, a new software tool. Front Microbiol [Internet]. Frontiers; 2019 [cited 2021 July 4];10. Available from: https://www.frontiersin.org/articles/10.3389/fmicb.2019.01391/full

13. Rennoll-Bankert KE, Garcia-Garcia JC, Sinclair SH, Dumler JS. Chromatin-bound bacterial effector ankyrin A recruits histone deacetylase 1 and modifies host gene expression. Cell Microbiol. 2015;17(11):1640–1652.

14. Lehman SS, Noriea NF, Aistleitner K, Clark TR, Dooley CA, Nair V, Kaur SJ, Rahman MS, Gillespie JJ, Azad AF, Hackstadt T. The rickettsial ankyrin repeat protein 2 is a Type IV secreted effector that associates with the endoplasmic reticulum. mBio. 2018 June 26;9(3):e00975–18. PMCID: PMC6020290

15. Lina TT, Farris T, Luo T, Mitra S, Zhu B, McBride JW. Hacker within! *Ehrlichia chaffeensis* effector driven phagocyte reprogramming strategy. Front Cell Infect Microbiol [Internet]. Frontiers; 2016 [cited 2021 June 30];6. Available from: https://www.frontiersin.org/articles/10.3389/fcimb.2016.00058/full#B90

16. Gillespie JJ, Phan IQH, Driscoll TP, Guillotte ML, Lehman SS, Rennoll-Bankert KE, Subramanian S, Beier-Sexton M, Myler PJ, Rahman MS, Azad AF. The Rickettsia type IV secretion system: unrealized complexity mired by gene family expansion. Pathog Dis [Internet]. 2016 Aug 1 [cited 2019 Mar 18];74(6). Available from: https://academic.oup.com/femspd/article/74/6/ftw058/2197957

17. IJdo JW, Carlson AC, Kennedy EL. *Anaplasma phagocytophilum* AnkA is tyrosine-phosphorylated at EPIYA motifs and recruits SHP-1 during early infection. Cell Microbiol. 2007;9(5):1284–1296.

18. Ramabu SS, Schneider DA, Brayton KA, Ueti MW, Graça T, Futse JE, Noh SM, Baszler TV, Palmer GH. Expression of *Anaplasma marginale* ankyrin repeat-containing proteins during infection of the mammalian host and tick vector. Infect Immun. 2011 July;79(7):2847–2855. PMCID: PMC3191954

19. Rikihisa Y, Lin M. *Anaplasma phagocytophilum* and *Ehrlichia chaffeensis* type IV secretion and Ank proteins. Curr Opin Microbiol. 2010 Feb;13(1):59–66.

20. Yan Q, Lin M, Huang W, Teymournejad O, Johnson JM, Hays FA, Liang Z, Li G, Rikihisa Y. *Ehrlichia* type IV secretion system effector Etf-2 binds to active RAB5 and delays endosome maturation. Proc Natl Acad Sci. Proceedings of the National Academy of Sciences; 2018 Sept 18;115(38):E8977–E8986.

21. Niu H, Kozjak-Pavlovic V, Rudel T, Rikihisa Y. *Anaplasma phagocytophilum* Ats-1 is imported into host cell mitochondria and interferes with apoptosis induction. PLOS Pathog. Public Library of Science; 2010 Feb 19;6(2):e1000774.

22. Wang L, Lin M, Hou L, Rikihisa Y. *Anaplasma phagocytophilum* effector EgeA facilitates infection by hijacking TANGO1 and SCFD1 from ER–Golgi exit sites to pathogen-occupied inclusions. Proc Natl Acad Sci. Proc Natl Acad Sci; 2024 Aug 13;121(33):e2405209121.

23. Gillespie JJ, Brayton KA, Williams KP, Diaz MAQ, Brown WC, Azad AF, Sobral BW. phylogenomics reveals a diverse Rickettsiales Type IV Secretion System. Infect Immun. 2010 May 1;78(5):1809–1823. PMID: 20176788

24. Garcia-Garcia JC, Rennoll-Bankert KE, Pelly S, Milstone AM, Dumler JS. Silencing of host cell CYBB gene expression by the nuclear effector AnkA of the intracellular pathogen *Anaplasma phagocytophilum*. Infect Immun. 2009 June 1;77(6):2385–2391. PMID: 19307214

25. Lin M, Liu H, Xiong Q, Niu H, Cheng Z, Yamamoto A, Rikihisa Y. *Ehrlichia* secretes Etf-1 to induce autophagy and capture nutrients for its growth through RAB5 and class III phosphatidylinositol 3-kinase. Autophagy. 2016 Nov;12(11):2145–2166. PMCID: PMC5103349

26. Biggs HM, Behravesh CB, Bradley KK, Dahlgren FS, Drexler NA, Dumler JS, Folk SM, Kato CY, Lash RR, Levin ML, Massung RF, Nadelman RB, Nicholson WL, Paddock CD, Pritt BS, Traeger MS. Diagnosis and management of tickborne rickettsial diseases: rocky mountain spotted fever and other spotted fever group rickettsioses, ehrlichioses, and anaplasmosis — United States: A Practical Guide for Health Care and Public Health Professionals. MMWR Recomm Rep. 2016 May 13;65(2):1–44.

27. Kumar S, Stecher G, Suleski M, Hedges SB. TimeTree: A resource for timelines, timetrees, and divergence times. Mol Biol Evol. 2017 July 1;34(7):1812–1819.

28. Epidemiology and Statistics | Anaplasmosis | CDC [Internet]. 2019 [cited 2019 Mar 13]. Available from: https://www.cdc.gov/anaplasmosis/stats/index.html

29. Lejal E, Moutailler S, Šimo L, Vayssier-Taussat M, Pollet T. Tick-borne pathogen detection in midgut and salivary glands of adult Ixodes ricinus. Parasit Vectors. 2019 Apr 2;12(1):152.

30. Ueti MW, Reagan JO, Knowles DP, Scoles GA, Shkap V, Palmer GH. Identification of midgut and salivary glands as specific and distinct barriers to efficient tick-borne transmission of *Anaplasma marginale*. Infect Immun. 2007 June;75(6):2959–2964. PMCID: PMC1932854

31. Park JM, Oliva Chávez AS, Shaw DK. Ticks: More than just a pathogen delivery service. Front Cell Infect Microbiol [Internet]. 2021 [cited 2022 Mar 29];11. Available from: https://www.frontiersin.org/article/10.3389/fcimb.2021.739419

32. Nelson CM, Herron MJ, Wang XR, Baldridge GD, Oliver JD, Munderloh UG. Global Transcription profiles of *Anaplasma phagocytophilum* at key stages of infection in tick and human cell lines and granulocytes. Front Vet Sci. 2020;7:111.

33. Nelson CM, Herron MJ, Felsheim RF, Schloeder BR, Grindle SM, Chavez AO, Kurtti TJ, Munderloh UG. Whole genome transcription profiling of *Anaplasma phagocytophilum* in human and tick host cells by tiling array analysis. BMC Genomics. 2008 Dec;9(1):1–16.

34. O’Conor MC, Herron MJ, Nelson CM, Barbet AF, Crosby FL, Burkhardt NY, Price LD, Brayton KA, Kurtti TJ, Munderloh UG. Biostatistical prediction of genes essential for growth of *Anaplasma phagocytophilum* in a human promyelocytic cell line using a random transposon mutant library. Pathog Dis. 2021 June 8;79(5):ftab029.

35. Oliva Chávez AS, Herron MJ, Nelson CM, Felsheim RF, Oliver JD, Burkhardt NY, Kurtti TJ, Munderloh UG. Mutational analysis of gene function in the Anaplasmataceae: Challenges and perspectives. Ticks Tick-Borne Dis. 2019 Feb 1;10(2):482–494.

36. Crosby FL, Munderloh UG, Nelson CM, Herron MJ, Lundgren AM, Xiao YP, Allred DR, Barbet AF. Disruption of VirB6 paralogs in *Anaplasma phagocytophilum* attenuates its growth. J Bacteriol. American Society for Microbiology; 202(23):e00301–20.

37. Oliva Chávez AS, Fairman JW, Felsheim RF, Nelson CM, Herron MJ, Higgins L, Burkhardt NY, Oliver JD, Markowski TW, Kurtti TJ, Edwards TE, Munderloh UG. An O-Methyltransferase Is required for infection of tick cells by *Anaplasma phagocytophilum*. PLoS Pathog. 2015 Nov 6;11(11):e1005248. PMCID: PMC4636158

38. Kuriakose JA, Miyashiro S, Luo T, Zhu B, McBride JW. *Ehrlichia chaffeensis* transcriptome in mammalian and arthropod hosts reveals differential gene expression and post transcriptional regulation. PLoS ONE. 2011 Sept 6;6(9):e24136. PMCID: PMC3167834

39. Villar M, Ayllón N, Alberdi P, Moreno A, Moreno M, Tobes R, Mateos-Hernández L, Weisheit S, Bell-Sakyi L, de la Fuente J. Integrated metabolomics, transcriptomics and proteomics identifies metabolic pathways affected by *Anaplasma phagocytophilum* infection in tick cells. Mol Cell Proteomics. 2015 Dec 1;14(12):3154–3172. PMCID: PMC4762615

40. Narra HP, Sahni A, Alsing J, Schroeder CLC, Golovko G, Nia AM, Fofanov Y, Khanipov K, Sahni SK. Comparative transcriptomic analysis of *Rickettsia conorii* during in vitro infection of human and tick host cells. BMC Genomics. 2020 Sept 25;21(1):665.

41. Riley SP, Pruneau L, Martinez JJ. Evaluation of changes to the *Rickettsia rickettsii* transcriptome during mammalian infection. PLoS ONE. 2017 Aug 23;12(8):e0182290. PMCID: PMC5568294

42. Mastronunzio JE, Kurscheid S, Fikrig E. Postgenomic analyses reveal development of infectious *Anaplasma phagocytophilum* during transmission from ticks to mice. J Bacteriol. 2012 May 1;194(9):2238–2247. PMID: 22389475

43. Barbet AF, Agnes JT, Moreland AL, Lundgren AM, Alleman AR, Noh SM, Brayton KA, Munderloh UG, Palmer GH. Identification of functional promoters in the *msp2* expression loci of *Anaplasma marginale* and *Anaplasma phagocytophilum*. Gene. 2005 June 20;353(1):89–97.

44. Lin Q, Rikihisa Y, Ohashi N, Zhi N. Mechanisms of variable p44 expression by *Anaplasma phagocytophilum*. Infect Immun. 2003 Oct 1;71(10):5650–5661. PMID: 14500485

45. Mirdita M, Schütze K, Moriwaki Y, Heo L, Ovchinnikov S, Steinegger M. ColabFold: making protein folding accessible to all. Nat Methods. Nature Publishing Group; 2022 June;19(6):679–682.

46. Holm L, Laiho A, Törönen P, Salgado M. DALI shines a light on remote homologs: One hundred discoveries. Protein Sci. 2023;32(1):e4519.

47. Rosenberg OS, Dovey C, Tempesta M, Robbins RA, Finer-Moore JS, Stroud RM, Cox JS. EspR, a key regulator of *Mycobacterium tuberculosis* virulence, adopts a unique dimeric structure among helix-turn-helix proteins. Proc Natl Acad Sci. Proc Natl Acad Sci; 2011 Aug 16;108(33):13450–13455.

48. Wang X, Kikuchi T, Rikihisa Y. Proteomic identification of a novel *Anaplasma phagocytophilum* DNA binding protein that regulates a putative transcription factor. J Bacteriol. 2007 July 1;189(13):4880–4886. PMID: 17483233

49. Wang X, Cheng Z, Zhang C, Kikuchi T, Rikihisa Y. *Anaplasma phagocytophilum* p44 mRNA expression is differentially regulated in mammalian and tick host cells: Involvement of the DNA binding protein ApxR. J Bacteriol. 2007 Dec 1;189(23):8651–8659. PMID: 17905983

50. Graça T, Ku PS, Silva MG, Turse JE, Hammac GK, Brown WC, Palmer GH, Brayton KA. Segmental variation in a duplicated *msp2* pseudogene generates *Anaplasma marginale* Antigenic Variants. Roy CR, editor. Infect Immun [Internet]. 2018 Nov 19 [cited 2019 Feb 14];87(2). Available from: http://iai.asm.org/lookup/doi/10.1128/IAI.00727-18

51. Graça T, Paradiso L, Broschat SL, Noh SM, Palmer GH. Primary structural variation in *Anaplasma marginale* Msp2 efficiently generates immune escape variants. Roy CR, editor. Infect Immun. 2015 Nov;83(11):4178–4184.

52. Lin Q, Zhang C, Rikihisa Y. Analysis of involvement of the RecF pathway in p44 recombination in *Anaplasma phagocytophilum* and in *Escherichia coli* by using a plasmid carrying the p44 expression and p44 donor loci. Infect Immun. 2006 Apr;74(4):2052–2062. PMCID: PMC1418890

53. Noroy C, Lefrançois T, Meyer DF. Searching algorithm for Type IV effector proteins (S4TE) 2.0: Improved tools for Type IV effector prediction, analysis and comparison in proteobacteria. PLoS Comput Biol. 2019 Mar;15(3):e1006847. PMCID: PMC6448907

54. Vondrak CJ, Sit B, Suwanbongkot C, Macaluso KR, Lamason RL. A conserved interaction between the effector Sca4 and host clathrin suggests additional contributions for Sca4 during rickettsial infection. Infect Immun. American Society for Microbiology; 2024 Nov 13;92(12):e00267–24.

55. Duan N, Ma X, Cui H, Wang Z, Chai Z, Yan J, Li X, Feng Y, Cao Y, Jin Y, Bai F, Wu W, Rikihisa Y, Cheng Z. Insights into the mechanism regulating the differential expression of the P28-OMP outer membrane proteins in obligatory intracellular pathogen *Ehrlichia chaffeensis*. Emerg Microbes Infect. 2021 Dec;10(1):461–471. PMCID: PMC7971322

56. Liu H, Knox CA, Jakkula LUMR, Wang Y, Peddireddi L, Ganta RR. Evaluating EcxR for Its possible role in *Ehrlichia chaffeensis* gene regulation. Int J Mol Sci. 2022 Oct 22;23(21):12719. PMCID: PMC9657007

57. Ge Y, Rikihisa Y. Identification of novel surface proteins of *Anaplasma phagocytophilum* by affinity purification and proteomics. J Bacteriol. American Society for Microbiology; 2007 Nov;189(21):7819–7828.

58. Villar M, Ayllón N, Kocan KM, Bonzón-Kulichenko E, Alberdi P, Blouin EF, Weisheit S, Mateos-Hernández L, Cabezas-Cruz A, Bell-Sakyi L, Vancová M, Bílý T, Meyer DF, Sterba J, Contreras M, Rudenko N, Grubhoffer L, Vázquez J, Fuente J de la. Identification and characterization of *Anaplasma phagocytophilum* proteins involved in infection of the tick vector, Ixodes scapularis. PLOS ONE. Public Library of Science; 2015 Sept 4;10(9):e0137237.

59. Contreras M, Alberdi P, Mateos-Hernández L, Fernández de Mera IG, García-Pérez AL, Vancová M, Villar M, Ayllón N, Cabezas-Cruz A, Valdés JJ, Stuen S, Gortazar C, de la Fuente J. *Anaplasma phagocytophilum* MSP4 and HSP70 proteins are involved in interactions with host cells during pathogen infection. Front Cell Infect Microbiol. 2017 July 5;7:307. PMCID: PMC5496961

60. Seidman D, Ojogun N, Walker NJ, Mastronunzio J, Kahlon A, Hebert KS, Karandashova S, Miller DP, Tegels BK, Marconi RT, Fikrig E, Borjesson DL, Carlyon JA. A*naplasma phagocytophilum* surface protein AipA mediates invasion of mammalian host cells. Cell Microbiol. 2014 Aug;16(8):1133–1145. PMCID: PMC4115035

61. Tang H, Zhu J, Wu S, Niu H. Identification and characterization of an actin filament-associated *Anaplasma phagocytophilum* protein. Microb Pathog. 2020 Oct 1;147:104439.

62. Zhu J, He M, Xu W, Li Y, Huang R, Wu S, Niu H. Development of TEM-1 β-lactamase based protein translocation assay for identification of *Anaplasma phagocytophilum* type IV secretion system effector proteins. Sci Rep. Nature Publishing Group; 2019 Mar 12;9(1):4235.

63. Sinclair SHG, Garcia-Garcia JC, Dumler JS. Bioinformatic and mass spectrometry identification of *Anaplasma phagocytophilum* proteins translocated into host cell nuclei. Front Microbiol. 2015 Feb 6;6:55. PMCID: PMC4319465

64. Cheng Z, Wang X, Rikihisa Y. Regulation of type IV Secretion apparatus genes during *Ehrlichia chaffeensis* intracellular development by a previously unidentified protein. J Bacteriol. American Society for Microbiology; 2008 Mar 15;190(6):2096–2105.

65. Shaw DK, Wang X, Brown LJ, Chávez ASO, Reif KE, Smith AA, Scott AJ, McClure EE, Boradia VM, Hammond HL, Sundberg EJ, Snyder GA, Liu L, DePonte K, Villar M, Ueti MW, de la Fuente J, Ernst RK, Pal U, Fikrig E, Pedra JHF. Infection-derived lipids elicit an immune deficiency circuit in arthropods. Nat Commun. 2017 14;8:14401. PMCID: PMC5316886

66. Chen G, Wang X, Severo MS, Sakhon OS, Sohail M, Brown LJ, Sircar M, Snyder GA, Sundberg EJ, Ulland TK, Olivier AK, Andersen JF, Zhou Y, Shi GP, Sutterwala FS, Kotsyfakis M, Pedra JHF. The tick salivary protein Sialostatin L2 Inhibits Caspase-1-mediated inflammation during *Anaplasma phagocytophilum* Infection. Infect Immun. 2014 June;82(6):2553–2564. PMCID: PMC4019176

67. Oliver JD, Chávez ASO, Felsheim RF, Kurtti TJ, Munderloh UG. An *Ixodes scapularis* cell line with a predominantly neuron-like phenotype. Exp Appl Acarol. 2015 July;66(3):427–442. PMCID: PMC4449809

68. Munderloh UG, Jauron SD, Fingerle V, Leitritz L, Hayes SF, Hautman JM, Nelson CM, Huberty BW, Kurtti TJ, Ahlstrand GG, Greig B, Mellencamp MA, Goodman JL. Invasion and intracellular development of the human granulocytic Ehrlichiosis agent in tick cell culture. J Clin Microbiol. American Society for Microbiology; 1999 Aug;37(8):2518–2524.

69. Sidak-Loftis LC, Rosche KL, Pence N, Ujczo JK, Hurtado J, Fisk EA, Goodman AG, Noh SM, Peters JW, Shaw DK. The unfolded-protein response triggers the arthropod immune deficiency pathway. mBio. American Society for Microbiology; 2022 July 18;0(0):e00703–22.

70. Sellegounder D, Liu Y, Wibisono P, Chen CH, Leap D, Sun J. Neuronal GPCR NPR-8 regulates *C. elegans* defense against pathogen infection. Sci Adv. American Association for the Advancement of Science; 2019 Nov 20;5(11):eaaw4717.

71. Kim D, Langmead B, Salzberg SL. HISAT: a fast spliced aligner with low memory requirements. Nat Methods. Nature Publishing Group; 2015 Apr;12(4):357–360.

72. Love MI, Huber W, Anders S. Moderated estimation of fold change and dispersion for RNA-seq data with DESeq2. Genome Biol. 2014;15(12):550. PMCID: PMC4302049

73. Cadby IT, Ibrahim SA, Faulkner M, Lee DJ, Browning D, Busby SJ, Lovering AL, Stapleton MR, Green J, Cole JA. Regulation, sensory domains and roles of two *Desulfovibrio desulfuricans* ATCC27774 Crp family transcription factors, HcpR1 and HcpR2, in response to nitrosative stress. Mol Microbiol. 2016;102(6):1120–1137.

74. Emsley P, Lohkamp B, Scott WG, Cowtan K. Features and development of Coot. Acta Crystallogr D Biol Crystallogr. International Union of Crystallography; 2010 Apr 1;66(4):486–501.

75. Pettersen EF, Goddard TD, Huang CC, Meng EC, Couch GS, Croll TI, Morris JH, Ferrin TE. UCSF ChimeraX: Structure visualization for researchers, educators, and developers. Protein Sci. 2021;30(1):70–82.

76. Wen Y, Behiels E, Felix J, Elegheert J, Vergauwen B, Devreese B, Savvides SN. The bacterial antitoxin HipB establishes a ternary complex with operator DNA and phosphorylated toxin HipA to regulate bacterial persistence. Nucleic Acids Res. 2014 Sept 2;42(15):10134–10147.

77. Gangwar SP, Meena SR, Saxena AK. Comparison of four different crystal forms of the *Mycobacterium tuberculosis* ESX-1 secreted protein regulator EspR. Acta Crystallogr Sect F Struct Biol Commun. International Union of Crystallography; 2014 Apr 1;70(4):433–437.

